# Intramuscular Adipose Tissue Accumulation is a Key Determinant of Limb Function in Peripheral Artery Disease

**DOI:** 10.64898/2026.01.27.701833

**Authors:** Victoria R. Palzkill, Divyansha Moparthy, Qingping Yang, Jaewon Choi, Xinyue Liu, Kyoungrae Kim, Ambili B. Appu, Caroline G. Pass, Scott A. Berceli, Curt D. Sigmund, Salvatore T. Scali, Daniel Kopinke, Terence E. Ryan

**Affiliations:** Department of Applied Physiology and Kinesiology, University of Florida, Gainesville, FL, USA; Department of Surgery, University of Florida, Gainesville, FL, USA; Department of Pharmacology and Therapeutics, University of Florida, Gainesville, FL, USA; Myology Institute, University of Florida, Gainesville, FL, USA; Center for Exercise Science, University of Florida, Gainesville, FL, USA; North Florida/South Georgia Veterans Health System, Malcom Randall VA Medical Center, Gainesville, FL, 32608; Department of Physiology, Medical College of Wisconsin, Milwaukee, WI, USA

**Keywords:** limb ischemia, muscle, fibro-adipogenic progenitor, FAPs, stem cell

## Abstract

**Background:** Peripheral artery disease (PAD) and its severe form, chronic limb-threatening ischemia (CLTI), significantly impair blood flow to the lower extremities, affecting millions of adults globally. Intramuscular adipose tissue (IMAT) and fibrosis accumulation distinguish patients with CLTI from those with mild PAD, suggesting a role in CLTI pathobiology. However, the functional consequences of IMAT in CLTI remain unclear.

**Methods:** We compared gastrocnemius muscle samples from patients with PAD/CLTI, intermittent claudication, and non-PAD individuals. We analyzed bulk RNA sequencing, proteomic, lipidomic, and single-cell/nucleus RNA sequencing datasets. Additionally, we used murine models of hindlimb ischemia (HLI) with genetic manipulation of Pparγ, a key adipogenic transcription factor, specifically in fibro-adipogenic progenitor cells (FAPs), the cellular source of IMAT, to modulate IMAT formation and assessed the impact on limb function and pathology.

**Results:** Patients with CLTI exhibited significantly elevated expression of adipogenic genes and proteins in muscle specimens when compared to non-PAD controls. Murine models showed that increasing IMAT formation significantly worsened ischemic limb muscle strength and work output. In contrast, preventing IMAT formation significantly improved ischemic limb muscle strength and work output. These findings were consistent across both male and female mice, although females had greater tendency to form IMAT compared with male mice.

**Conclusions:** IMAT accumulation is a key determinant of limb function in PAD/CLTI. Our studies demonstrate that targeting IMAT formation could improve limb function in mice with experimental PAD. Together, these findings suggest that developing strategies to limit or reduce IMAT may improve limb function and walking performance in patients with PAD/CLTI, providing a novel therapeutic avenue to address a critical unmet need.

**CLINICAL PERSPECTIVE:** *What is new?:*

- Intramuscular adipose tissue accumulation (IMAT) distinguishes patients with chronic limb-threatening ischemia from those with milder peripheral artery disease or those without PAD and directly impairs ischemic limb muscle function.
- Genetic gain- and loss-of-function mouse models demonstrate that increasing IMAT worsens, while preventing IMAT formation improves, ischemic limb strength and performance independent of perfusion.
- Adipogenic signatures in human calf muscle negatively correlates with muscle strength and disease severity, identifying IMAT as a functional biomarker and modifiable target in PAD/CLTI.

*What are the clinical implications?:*

- IMAT accumulation represents an underappreciated, non-vascular mechanism contributing to leg dysfunction in PAD/CLTI.
- Therapies aimed at limiting or reversing IMAT formation may improve leg strength and walking performance in patients with PAD/CLTI, addressing a critical unmet clinical need.
- Identifying and targeting cellular pathways regulating IMAT formation from fibro-adipogenic progenitors may complement vascular interventions to enhance functional recovery after revascularization.

## INTRODUCTION

Peripheral artery disease (PAD) is a chronic atherosclerotic condition that impairs blood flow to the lower extremities, affecting approximately 8–12 million adults in the United States and >200 million globally, making it a major cause of cardiovascular morbidity and mortality^1-3^. Its most severe form, chronic limb-threatening ischemia (CLTI), is characterized by rest pain, non-healing ulcers, and gangrene, and is associated with higher risk of limb amputation and death. Current standard of care for PAD/CLTI involves re-establishing blood flow to the ischemic tissue, yet endovascular and revascularization procedures have high failure rates^4-6^. In addition, numerous cell- and gene-based vascular therapies had sub-optimal efficacy in clinical trials^7-10^. While the failure of vascular-focused therapies is multifactorial, there is a limited understanding of how non-vascular cells, and their corresponding cellular interactions contribute to PAD/CLTI pathobiology. This void in knowledge is a significant barrier to developing effective therapies to improve limb function in PAD/CLTI.

The accumulation of non-vascular tissues such as intramuscular fat (IMAT) and fibrosis distinguishes mild PAD patients with claudication from those with severe PAD/CLTI^11-13^, implicating IMAT and fibrosis as possible mechanisms contributing to the CLTI disease manifestation. However, the functional consequences of IMAT and/or fibrosis in CLTI are unknown. In patients with PAD/CLTI, limb function is a strong predictor of morbidity and mortality^14-17^. Thus, we hypothesized that the replacement of muscle fibers with IMAT and/or fibrotic tissue are likely contributors to the weakness and poor muscle function observed in patients with PAD/CLTI^15,18-21^.

Here, we report that the accumulation of IMAT, which is driven by the differentiation of fibro/adipogenic progenitors (FAPs) into adipocytes^22-27^ is a clinically important pathological hallmark of PAD/CLTI and adipogenic gene expression in the gastrocnemius muscle negatively associates with ischemic calf muscle function. Through comprehensive *in vivo* genetic gain- and loss-of-function and phenotyping strategies, we demonstrate that increasing IMAT formation worsens ischemic limb function, whereas preventing IMAT formation significantly improved ischemic limb function. Collectively, these studies demonstrate that IMAT accumulation is a key determinant of limb function in PAD/CLTI and that developing strategies to alter IMAT levels hold promise for improving leg function.

## MATERIALS AND METHODS

Detailed descriptions of the materials and experimental procedures can be found in the Expanded Material and Methods section of the **Supplemental Material**. The data supporting the conclusions of this study are available from the corresponding author. A Major Resources Table is available in the **Supplemental Material**.

### Human Studies and Datasets

Gastrocnemius muscle specimens were collected from PAD/CLTI and non-PAD patients via percutaneous muscle biopsy using sterile procedures as previously described ^28,29^ and used to generate total RNA for gene expression analysis (non-PAD n=18, CLTI n=38). CLTI was confirmed by a Rutherford Stage of 4-5 consisting of ischemic rest pain for more than two weeks. This study was approved by the institutional review boards at the University of Florida and the Malcom Randall VA Medical Center (Protocol IRB201801553). All study procedures were carried out according to the Declaration of Helsinki and participants were fully informed about the research and informed consent was obtained. A detailed description of the patient cohort has been previously published^30^ and key physical and clinical characteristics can be found in Supplemental Table 2. We also analyzed several publicly available and previously published datasets involving human patients with PAD/CLTI. Bulk RNA sequencing data from the gastrocnemius muscle of non-PAD, patients with intermittent claudication, and patients with CLTI undergoing limb amputation was obtained from our lab^28^ (GSE120642). Proteomic^29^ and lipidomic^31^ data from non-PAD volunteers and patients with CLTI undergoing limb amputation was obtained from our lab (jPost Accession # PXD021849 and Metabolomics Workbench Study ID’s ST001615, ST001616, ST001617). Single-cell RNA sequencing data from patients with CLTI undergoing limb amputation was obtained from Southerland *et al*.^32^ (GSE227077). Single-nucleus RNA sequencing data from non-PAD volunteers and patients with PAD/CLTI was obtained from our lab^30,33^ (GSE233882 and GSE284482). Patients included in the single-nucleus RNA sequencing datasets had Rutherford stages 3-5, so they are described as PAD and not CLTI here forth.

### Animal Studies

All animal experiments adhered to the *Guide for the Care and Use of Laboratory Animals* from the Institute for Laboratory Animal Research, National Research Council, Washington, D.C., National Academy Press. All procedures were approved by the Institutional Animal Care and Use Committee of the University of Florida (Protocol 202300000490). Mice were housed in a temperature- (22°C) and light-controlled (12:12-h light-dark) room and maintained on standard chow (Envigo Teklad Global 18% Protein Rodent Diet 2918 irradiated pellet) with free access to food and water. BALB/cJ mice (Stock No. 000651) were obtained from Jackson Laboratories. To manipulate IMAT formation, we targeted the transcription factor *Pparγ*, a key regulator of adipogenesis, using a tamoxifen-inducible, FAP-specific *Pdgfrα*^*CreERT2*^ mouse (Jackson Laboratories, Stock No. 032770)^34^. *Pdgfrα*^*CreERT2*^ mice were bred with conditional transgenic mice expressing a human wild-type PPARG allele under control of the CAG promoter (CAG-LSL-PPARG; kindly provided by Dr. Curt Sigmund, Medical College of Wisconsin)^35^ to generate a FAP-specific, inducible overexpression of PPARγ model termed mFATGAIN. To block IMAT formation, crossed the *Pdgfrα*^*CreERT2*^ mice with floxed *Pparg* mice (B6.129-*Ppargtm2Rev*/J (PPARγ^loxP^, Jackson Laboratories Stock No. 004584) carrying an inducible EYFP reporter (B6.129X1-*Gt(ROSA)26Sortm1(EYFP)Cos*/J, Jackson Laboratories Stock No. 006148) which we termed mFATBLOCK^36^. All FAP-specific mouse lines were backcrossed to the 129S1/SvlmJ genetic background because C57BL/6J mice are resistant to limb ischemia and form little IMAT with non-ischemic injuries^37^. In all experiments, littermates lacking the Cre transgene were used as controls. All experiments involved male and female mice that were randomized to experimental groups so that both the surgeon and experimenters were blinded to the treatment groups and genotypes. Female mice underwent an ovariectomy two weeks prior to enrollment. All mice received tamoxifen (TRC, T006000; 200-250 mg/kg) dissolved in sunflower seed oil via oral gavage for three consecutive days to induce Cre recombination. Two weeks after tamoxifen administration, femoral artery ligation was performed to induce unilateral hindlimb ischemia (HLI)^38,39^. Perfusion recovery was assessed by laser Doppler perfusion imaging (Model LDI2-IR, Moor Instruments, Wilmington, DE). At either 7- or 28-days post-HLI, limb function was evaluated using a 6-minute limb function test^40^, and tissues were collected for histological analysis. Body composition and blood glucose were assessed after tamoxifen administration but prior to surgery, using EchoMRI and tail vein blood sampling, respectively.

### Statistical Analysis

Normality of data was tested with the Shapiro-Wilk test and inspection of QQ plots. Data are presented as mean ± standard error of the mean (SEM) unless otherwise indicated. For comparisons between two groups, Student’s t-tests were used when data were normally distributed. If normality could not be established, or sample size was < 6, the non-parametric Mann-Whitney test was applied. Two-way ANOVA was used to analyze the effects of multiple factors across experimental groups with Tukey’s post hoc testing for multiple comparisons when significant interactions were detected. Perfusion recovery over time was analyzed using mixed-effects models to account for repeated measures. Univariable and multivariable linear regression models were used to examine the relationship between limb function (total work and muscle strength) and other outcome variables (IMAT, total myofiber area, average myofiber CSA, paw perfusion, total capillary density, perfused capillary density). As the relationship between IMAT and limb function was not linear, IMAT values were log transformed for the regression analyses. Variance inflation factor was used to check for multicollinearity of linear regression models, and a variance inflation factor of <4 was used to define absence of multicollinearity. For all analyses, *P* < 0.05 was considered statistically significant. Regression analyses were performed in SPSS (version 29.0.2.0) and all other statistical testing was conducted using GraphPad Prism software (version 10.0).

## RESULTS

### IMAT is a hallmark CLTI muscle in both human patients and mice

Using CT scans, previous work has shown that the percentage of fat within the calf muscle increases with PAD severity^41^. In patients with CLTI, white adipocytes are easily visualized in gastrocnemius muscle sections (**Figure 1A**). Examination of a bulk RNA sequencing dataset from the gastrocnemius muscle comparing healthy adults (HA, n=15), patients with intermittent claudication (IC, mild form of PAD, n=20), and those with CLTI (n=16)^28^ revealed that adipogenic genes (*ADIPOQ, ADIRF, FABP4, CEBPA, PLIN1*, and *PPARG*) were significantly elevated specifically in patients with CLTI (**Figure 1B**). On a separate cohort of non-PAD volunteers (n=18) and patients with CLTI (n=38; participant characteristics can be found in the Supplemental Material)^30^, qPCR analysis also demonstrated significantly increased expression of *ADIPOQ, FABP4*, and *CEBPA* in CLTI muscles (**Figure 1C**). Analysis of our existing proteomics dataset (non-PAD n=9, CLTI n=11)^42^ revealed a higher abundance of several adipocyte-associated proteins such as PLIN1, ADIPOQ, and ADIRF in muscle from patients with CLTI when compared to non-PAD controls (**Figure 1D**). The accumulation of IMAT in limb muscle from patients with CLTI was also present in a lipidomic dataset (non-PAD n=8, CLTI n=12)^43^ (**Figure 1E**). To assess the adipogenic potential of the FAPs, the cellular origin of IMAT, we first examined a single-cell RNA-sequencing dataset from Southerland *et al*.^44^ which quantified RNA expression in 15,843 mononuclear cells isolated from ischemic (distal) and non-ischemic (proximal) muscle of patients with CLTI undergoing limb amputation (n=3). Here the FAPs displayed significantly higher expression of adipogenic genes *PPARG, CEBPA*, and *ANGPLT4* ^45,46^ in the ischemic (distal) samples compared to the non-ischemic (proximal) samples (**Figure 1F**), confirming that the CLTI microenvironment promotes a pro-adipogenic phenotype in resident FAPs. Next, we pooled single-nucleus RNA-sequencing data from our two published datasets^30,47^ comprising 72,902 nuclei from gastrocnemius biopsies of patients with PAD (n=32) and non-PAD controls (n=12). Consistent with the Southerland *et al*. dataset, FAPs from patients with PAD had higher expression of *PPARG* compared to FAPs from non-PAD controls (**Figure 1G**). Collectively, the consistent increase of adipogenesis-related genes/proteins in the limb muscle of patients with PAD/CLTI across numerous independent datasets obtained from different patient cohorts establish IMAT as a hallmark of CLTI. Next, we quantified the temporal development of IMAT in the ischemia susceptible BALB/cJ mouse, known to exhibit CLTI-like features including impaired limb perfusion, limb and muscle necrosis, and deficient angiogenesis following HLI ^40,48-50^. Consistent with patients, BALB/cJ mice displayed a rapid and substantial increase in the number of Perilipin^+^ adipocytes present in the ischemic hindlimb muscle (**Figure 1H**).

**Figure 1.**
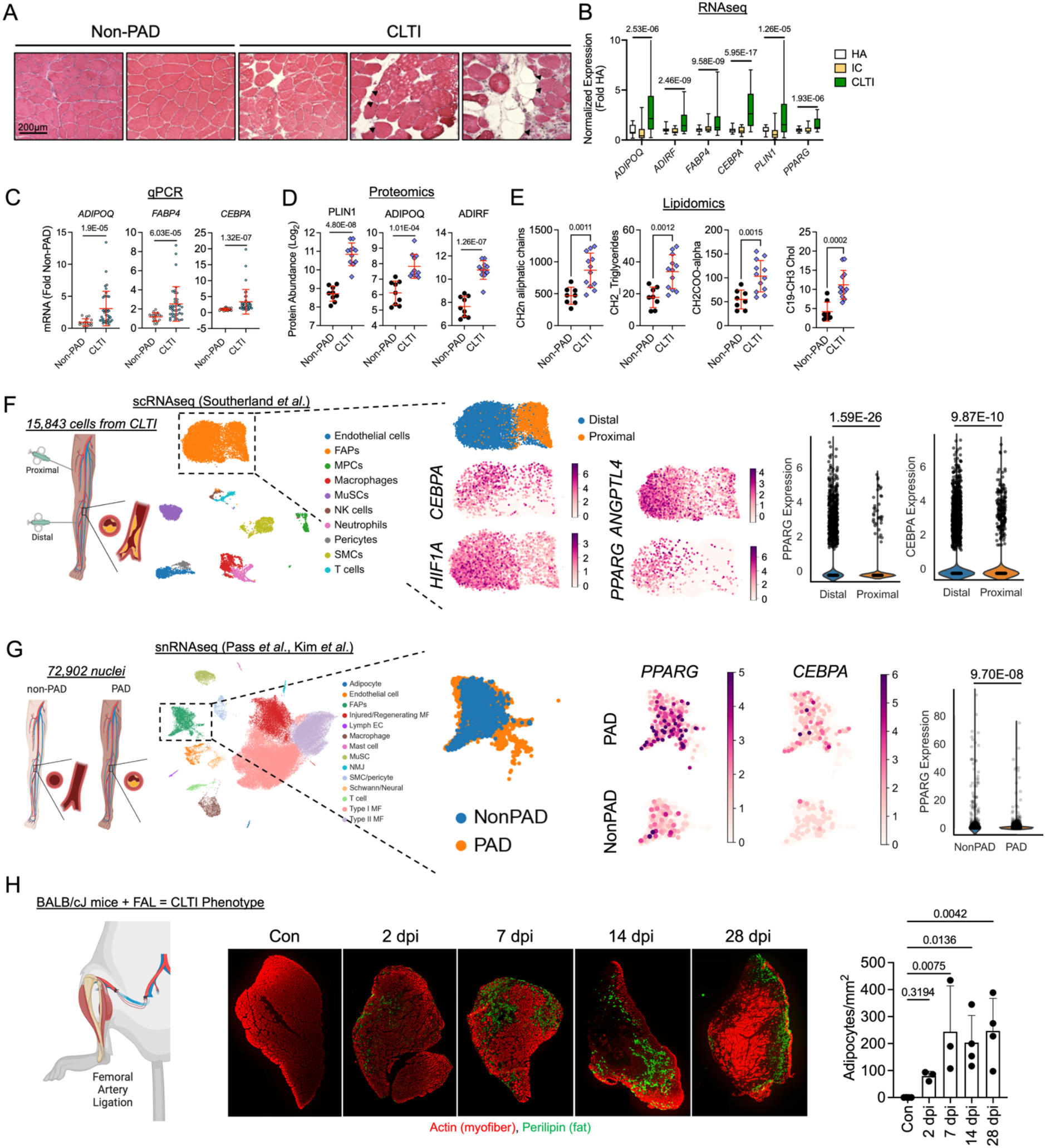
Accumulation of IMAT is a hallmark of human and murine PAD/CLTI. (A) Hematoxylin and eosin staining of gastrocnemius muscle from non-PAD and CLTI participants. Black triangles highlight adipocytes. (B) IMAT associated gene expression from an available RNA sequencing dataset (HA, healthy adults, n=15; IC, intermittent claudicants, n=20; CLTI, chronic limb threatening ischemic, n=16). (C) Relative mRNA expression of adipogenesis related genes (non-PAD n=18, CLTI n=38). (D) Normalized protein abundance (Log2) of IMAT-associated proteins (non-PAD n=9 patients, CLTI n=11 patients). (E) Lipidomic analysis of non-PAD and CLTI muscle specimens (non-PAD n=8 patients, CLTI n=12 patients). (F) scRNAseq analysis of FAPs from patients with CLTI including feature plots of FAPs and violin plots of expression for adipogenic genes. (G) snRNAseq analysis of PAD and Non-PAD muscles with feature plots and a violin plot for adipogenic gene expression. (H) Representative images of IMAT stained with perilipin and quantification over time in BALB/c mice subjected to femoral artery ligation (FAL) (control n= 3 mice, 2dpi n= 3 mice, 7dpi n=3 mice, 14 dpi n=4 mice, 28dpi n=4 mice). Error bars represent the standard deviation.

### FAP-specific overexpression of *Pparγ* does not induce IMAT in the absence of ischemia

Although IMAT accumulation is a consistent feature of PAD/CLTI in both human and murine limbs, its role in PAD/CLTI pathobiology is unknown. To address this gap in knowledge, we generated a genetic gain-of-function model using tamoxifen-inducible, FAP-specific overexpression of the human PPARγ transgene, termed mFATGAIN (**Figure 2A**). Following tamoxifen administration, qPCR analysis of FAPs revealed a >100-fold increase in human PPARγ expression compared to controls, which resulted in downregulation of the endogenous mouse Pparγ mRNA expression (**Figure 2B**). No human PPARγ expression was detected in the Cre^-^ controls. FAP-specific expression of the human PPARγ transgene significantly increased the expression of downstream endogenous adipogenic genes *Fabp4* and *Cebpa* (**Figure 2B**). Importantly, mFATGAIN mice did not exhibit differences in body weight, body composition, or blood glucose levels compared to controls (**Figure 2C**). In the absence of ischemia, mFATGAIN mice had normal gastrocnemius muscle weights (**Figure 2D**) and no differences in FAP abundance (**Figure 2E and Supplemental Figure 1**), myofiber area (**Figure 2F**), adipocyte number (**Figure 2G, H**), or total and perfused capillary densities (**Figure 2I**) when compared to littermate controls. These data demonstrate that inducible PPARγ overexpression in FAPs does not induce IMAT formation or alter overall muscle homeostasis in non-ischemic muscle.

**Figure 2.**
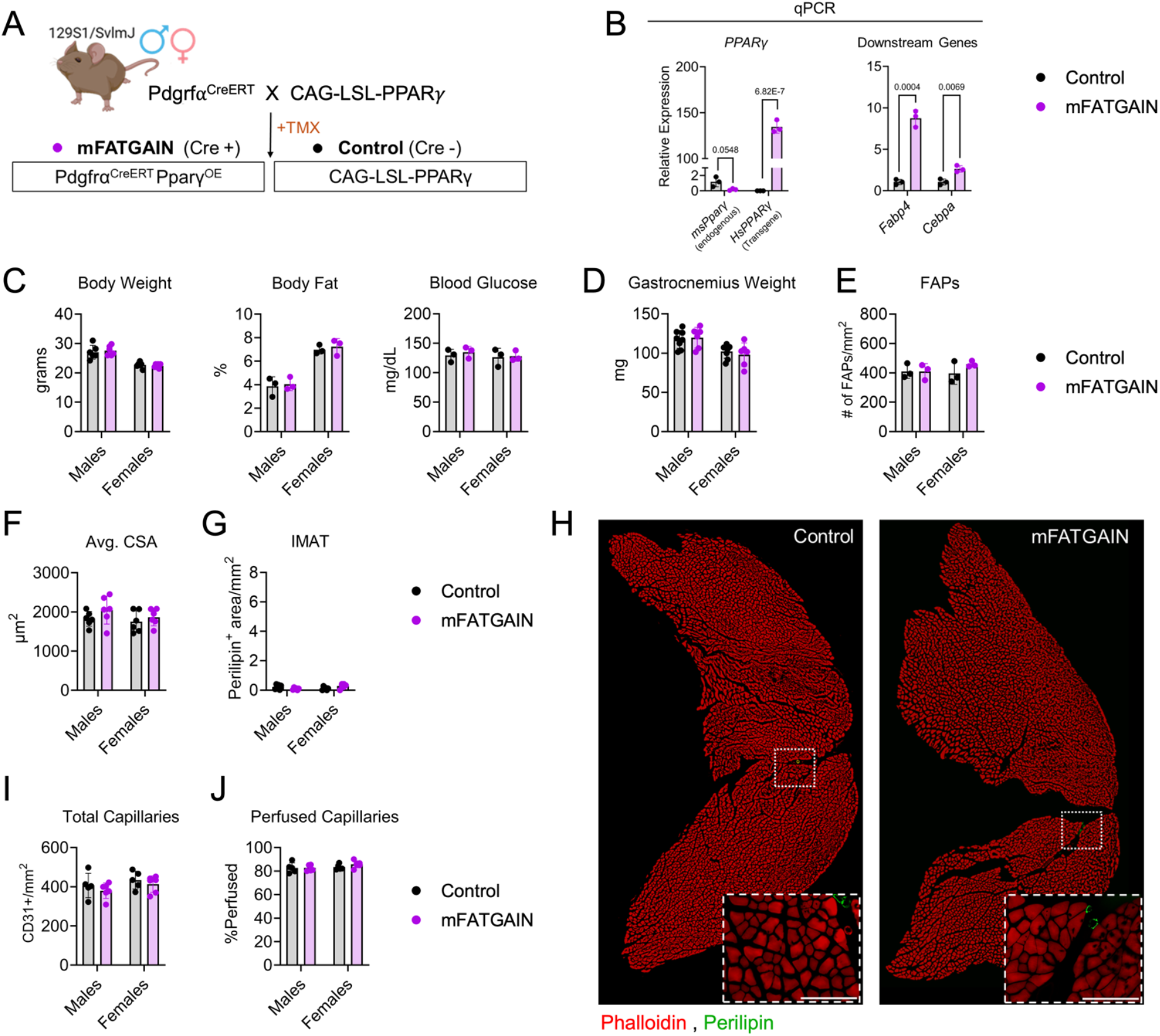
FAP-specific PPARγ overexpression does not alter homeostasis in non-ischemic muscle. (A) Schematic of tamoxifen-inducible, FAP-specific PPARγ overexpression mouse model (mFATGAIN) generated by crossing CAG-LSL-PPARG mice with Pdgfrα^CreERT2^ mice (B) Transgene activation and downstream adipogenic gene expression assessed by qPCR in FAPs isolated from hindlimb muscle (n= 3 biological replicates/group) and analyzed using a Mann-Whitney test. (C) Basic metabolic phenotyping via body weight (n=6/group/sex), body fat percentage, and blood glucose (n = 3/genotype/sex). (D) Wet weight of non-ischemic gastrocnemius (control n = 8/sex, mFATGAIN n = 6/sex). (E) Number of FAPs stained for PDGFRa per mm^2^ (n=3/group/sex). (F, G) Average myofiber CSA (F) and total IMAT area (G) quantified using Perilipin staining (n = 6/group/sex). (H) Representative phalloidin and Perilipin-stained images (20x magnification); scale bar = 200 μm. (I) Total capillaries probed with anti-CD31 antibody and normalized to area. (J) Perfused capillaries labeled with retro-orbitally injected isolectin and reported as a percentage of total capillaries (n=5/sex control, n=6/sex mFATGAIN). (C-J) Analyzed using two-way ANOVA. Data shown as mean ± SD.

### Overexpression of PPARγ increases IMAT formation in mice subjected to HLI

To assess the effects of FAP-specific PPARγ overexpression under ischemic conditions, mFATGAIN and littermate control mice were subjected to HLI and muscles were analyzed 7- and 28-days post-HLI (**Figure 3A**). Laser Doppler imaging showed no differences in paw perfusion recovery between groups in either sex (**Figure 3B**). Similarly, mFATGAIN and control mice had no differences in total or perfused capillary densities within the ischemic gastrocnemius, (**Figure 3C-E**). Perilipin staining of the ischemic gastrocnemius revealed a striking increase (∼5-17 fold) in IMAT levels of both male and female mFATGAIN mice at both timepoints when compared to littermate controls (**Figure 3G** and **Supplemental Figure 2**). However, the total FAP abundance was not different between groups, except for a modest reduction in male mFATGAIN mice at 28-days post-HLI (**Figure 3H**), indicating that expression of Ppary and does not cause any major shift in FAP numbers. Whereas the overall gastrocnemius muscle weight was not different between groups (**Figure 3I**), mFATGAIN mice had significantly smaller muscle fiber cross-sectional areas at both 7- and 28-days post-HLI (**Figure 3J**). At 7-days post-HLI, mFATGAIN mice had a greater abundance of myofibers expressing embryonic myosin heavy chain (Myh3) when compared to controls at (**Supplemental Figure 3)** indicating delayed myofiber regeneration. mFATGAIN mice also displayed increased abundance of leukocytes (CD45+), macrophages (CD68+) and neutrophils (Ly6G+) in the ischemic gastrocnemius muscle when compared to littermate controls (**Supplemental Figure 4A-D**). Despite a significant increase in immune cell abundance, only modest changes in cytokine concentrations in the whole muscle were observed (**Supplemental Figure 4E)**.

**Figure 3.**
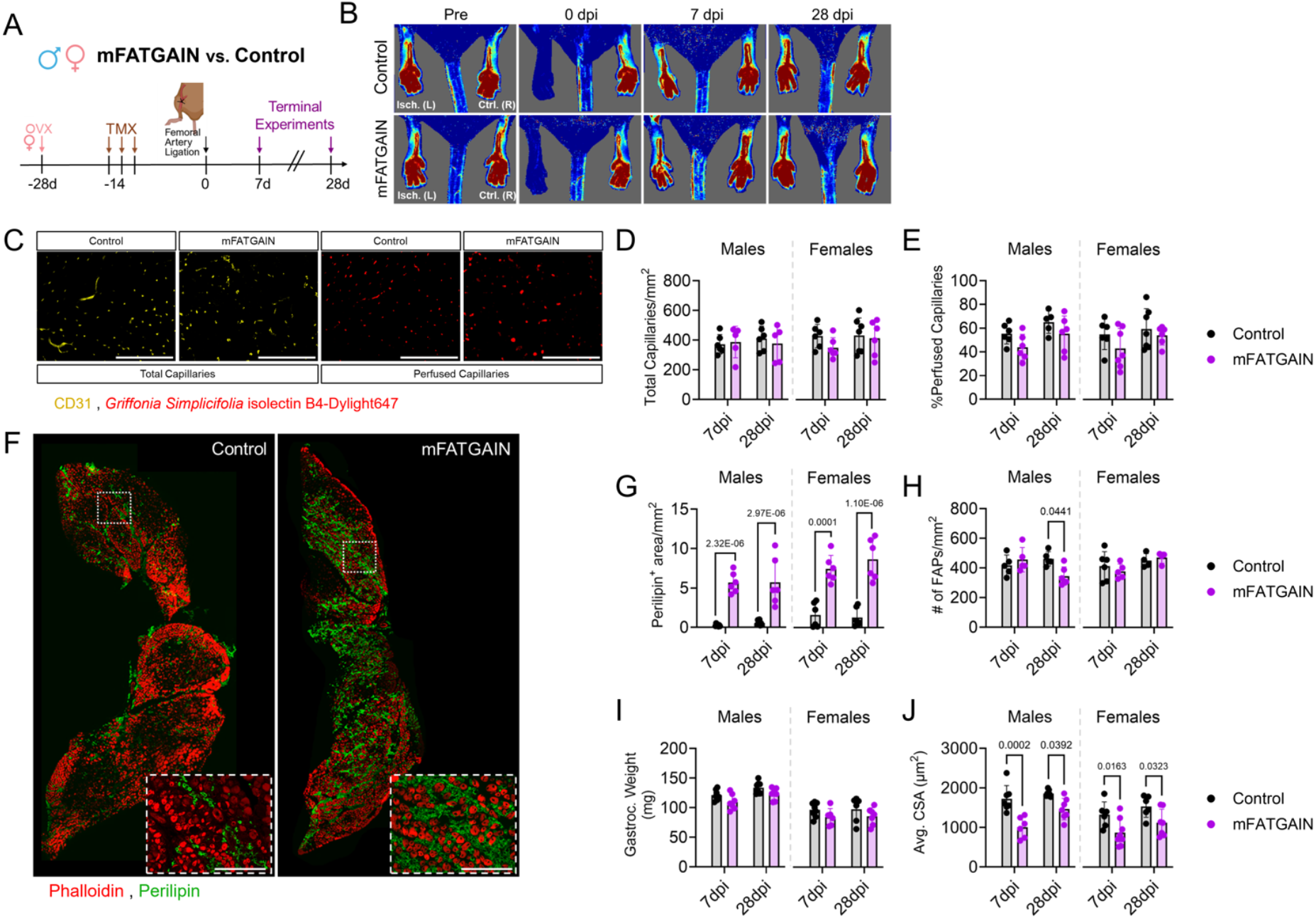
FAP-specific PPARγ overexpression increases IMAT formation following HLI. (A) Schematic of experimental design. (B) Resting paw perfusion measured with Laser Doppler imaging throughout recovery period (n=8/group/timepoint). (C–F, I) Ischemic gastrocnemius analysis at 7- and 28-days post-injury (dpi). (C) Representative images of total (CD31^+^) and perfused (isolectin^+^) capillaries and corresponding quantifications (D,E) in males (7dpi; controls n=6, mFATGAIN n=5. 28dpi; controls n=6, mFATGAIN n=5) and females (7dpi; n=6/genotype. 28dpi; controls n=7, mFATGAIN n=6). (F) Representative images of IMAT (Perilipin) and myofibers (Phalloidin). (G) Quantification of IMAT area in males (7dpi; control n = 7, mFATGAIN n = 6. 28dpi; control n = 8, mFATGAIN n = 6) and females (7dpi; n = 6/genotype. 28dpi; n = 6/genotype). (H) Number of FAPs quantified with PDGFRa staining and normalized to area in males (7dpi; controls n=5, mFATGAIN n=6. 28dpi; n=6/genotype) and females (7dpi; controls n=5, mFATGAIN n=6. 28dpi; n=6/genotype). (I) Muscle wet weight and (J) average myofiber CSA in males (7dpi; control n = 7, mFATGAIN n = 6. 28dpi; n = 7/genotype) and females (7dpi; n = 8/genotype. 28dpi; control n = 7, mFATGAIN n = 8). Representative immunofluorescent images at 20x magnification from female mice at 7dpi, scale bars = 200 μm. Data were analyzed using a two-way ANOVA with Tukey’s post hoc testing for multiple comparisons when significant interactions were detected. All data are presented as mean ± SD.

### Increased IMAT impairs muscle function in mice with HLI

To evaluate the functional consequences of increased IMAT, nerve-mediated *in-situ* contraction of the plantar flexor complex was performed. Muscle strength was evaluated using a series of isometric contractions across increasing stimulation frequencies (1,40, 80, 100 and 150Hz). Consistent with our hypothesis, mFATGAIN mice had significantly weaker muscles compared to littermate controls following HLI (**Figure 4A**). Quantification of specific force (absolute force normalized by muscle weight), an indicator or muscle quality, also demonstrated that mFATGAIN mice had significantly lower specific forces than their littermates (**Figure 4B**). mFATGAIN mice also has significantly lower muscle power output at 30% of an 80Hz contraction (**Figure 4C**). Because the increased IMAT formation in mFATGAIN mice did not alter the total muscle weight, we thought differences in specific force may be due to mFATGAIN mice having fewer myofilament proteins rather than intrinsic myofilament protein dysfunction. Quantification of the total area of the gastrocnemius muscle occupied by myofibers confirmed that mFATGAIN mice had less total myofiber area within the ischemic limb (**Supplemental Figure 5A, B**). Normalization of absolute forces to total myofiber areas revealed similar muscle quality between mFATGAIN and control mice at 28-days post-HLI (**Supplemental Figure 5C**). However, at 7-days post-HLI absolute forces normalized to total myofiber areas uncovered a deficit in muscle quality in mFATGAIN mice (**Supplemental Figure 5C**), suggesting that increases in IMAT delay functional recovery with ischemic injury without causing long-term damage to the contractile apparatus.

**Figure 4.**
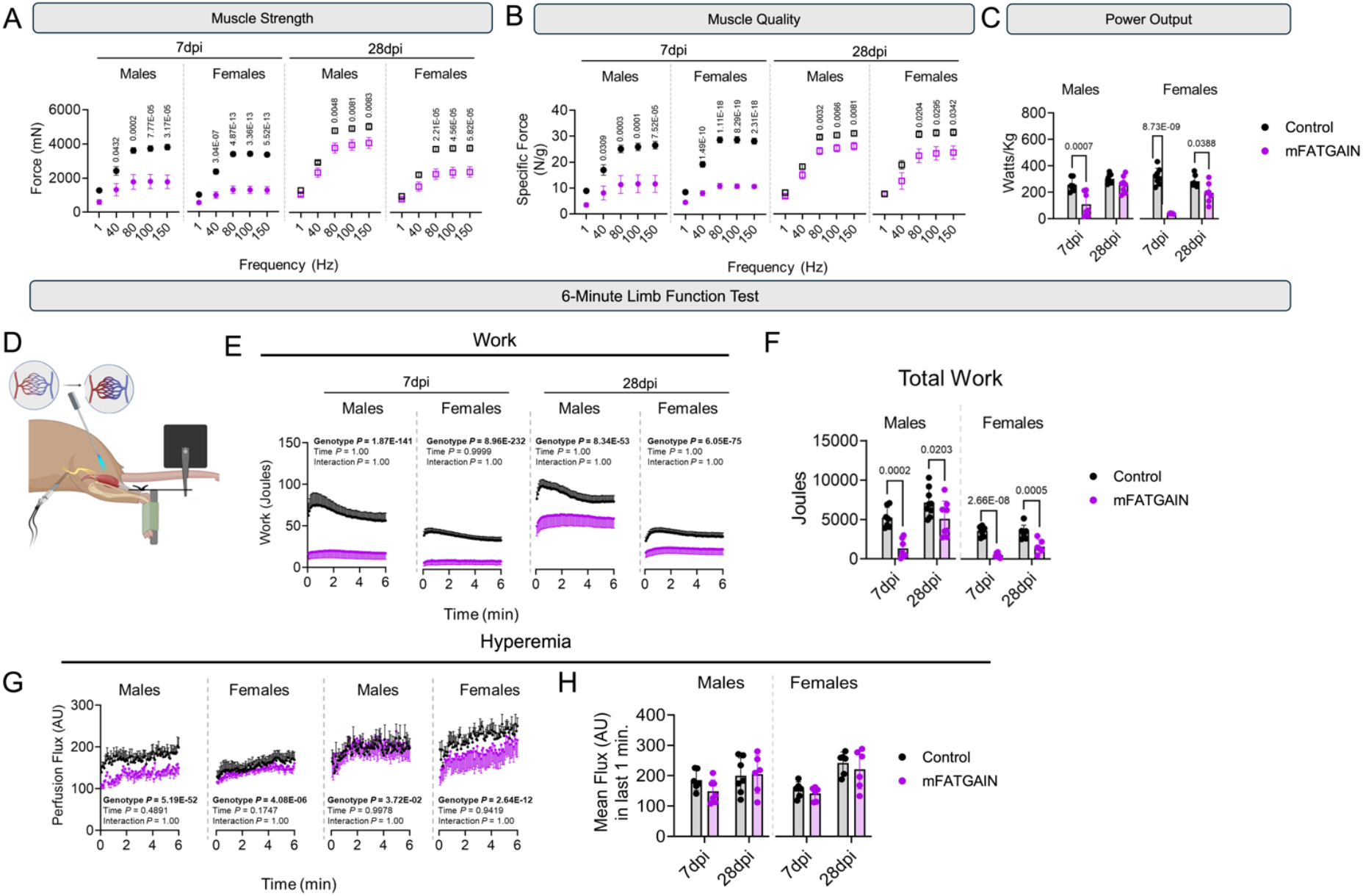
Increased IMAT formation impairs muscle function in mice with HLI. Tetanic contractions at increasing stimulation frequencies (1,40, 80, 100 and 150Hz) reported as absolute force (A), and (B) specific force normalized to muscle weight. (C) Power generation at 30% of 80Hz tetanic force. (D) Schematic of 6-minute limb function test. (E) Work performed across the 6-minute limb function test and (F) total work performed during the test. (A-F) Males (7dpi; n=7 mice control, n= 8 mice mFATGAIN. 28dpi; n=8 mice/genotype) and females (7dpi; n=8 mice/genotype 28dpi; n=7 mice/genotype). (G) Laser Doppler perfusion flux in the gastrocnemius across the test and (H) average flux during the last one minute in males (7dpi; control n=5 mice, mFATGAIN n= 7. 28dpi; control n=7 mice, mFATGAIN n= 6 mice) and females (7dpi; n=6 mice/genotype 28dpi; n= 6 mice/genotype). Data were analyzed using a two-way-ANOVA with Tukey’s post hoc testing when appropriate. Error bars represent ± SD.

Next, we employed a 6-minute limb function test to comprehensively assess ischemic muscular performance^40^. This test uses repeated isotonic (shortening) contractions with continuous laser Doppler flowmetry to allow for quantification of muscle work output and hyperemia (**Figure 4D**). Across the 6-min limb function test, both male and female mFATGAIN mice display significantly lower muscular work output (**Figure 4E** and **Supplemental Table 3**) resulting in a decreased total work performed (a measure akin to 6-min walk distance for human PAD trials as shown in **Figure 4F**. Quantitatively, at 7-days post-HLI, total work performed for mFATGAIN mice was 1163 ± 1190 and 619 ± 460 Joules for males and females respectively. In contrast, WT littermates performed 5094 ± 2244 (males) and 3581 ± 585 (females) Joules of total work at 7-days post-HLI. Notably, gastrocnemius perfusion was not different between mFATGAIN and control mice during the 6-min limb function test (**Figure 4G, H**), suggesting that limb perfusion is independent of IMAT.

### FAP-specific Pparγ deletion diminishes IMAT formation in the ischemic limb

Next, we sought to test if FAP-specific deletion of Pparγ would suppress IMAT formation following HLI. To accomplish this, we generated a tamoxifen-inducible, FAP-specific Pparγ knockout model, termed mFATBLOCK (**Figure 5A**). With this model, we achieve recombination in ∼88% of FAPs, evidenced by positive expression of the EYFP reporter (**Supplemental Figure 6A**). qPCR of isolated FAPs confirmed a robust reduction in expression of *Pparg* and its downstream targets *Cebpa* and *Fabp4* in mFATBLOCK mice compared to littermate controls (**Figure 5B**). Importantly, no differences in body weight, body fat, or blood glucose levels were observed (**Figure 5C**). Moreover, without ischemia, deletion of Pparγ in FAPs did not significantly affect FAP abundance, IMAT levels, muscle mass, myofiber area, or capillary density in the gastrocnemius muscle (**Supplemental Figure 6B-F**).

**Figure 5.**
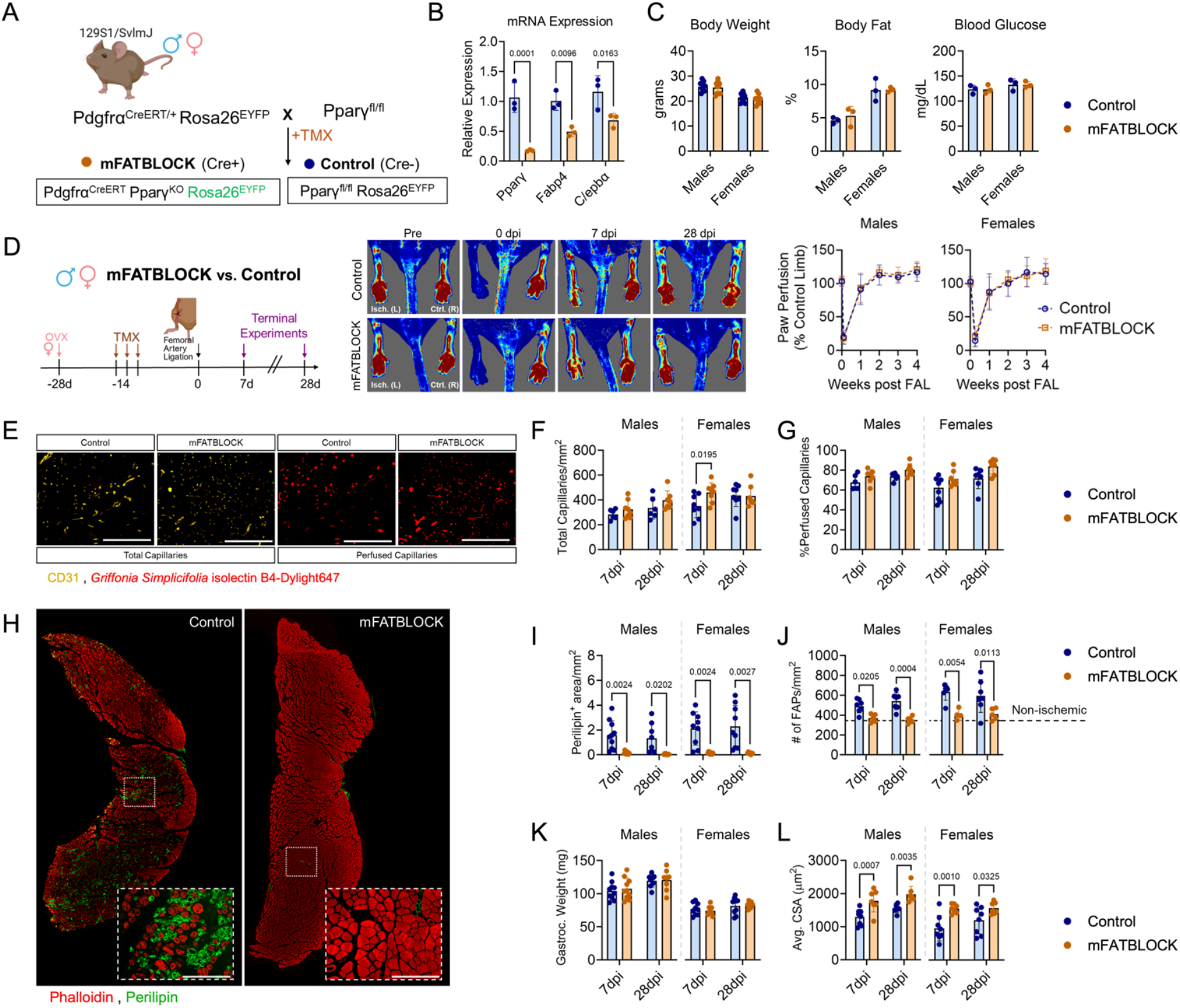
mFATBLOCK mice are protected against ischemia induced IMAT formation. (A) Schematic of tamoxifen-inducible, FAP-specific *Pparγ* knockout mouse model (mFATBLOCK). (B) *Pparγ* deletion and reduced adipogenic gene expression in FAPs from hindlimb muscle confirmed by qPCR and analyzed using a Mann-Whitney test (n = 3 biological replicates/group). (C) Body weight (n = 8/genotype/sex), body fat percentage, and blood glucose (n = 3/genotype/sex). (D) Paw perfusion recovery assessed by Laser Doppler imaging and analyzed using a mixed-effects model (n = 9/group/timepoint). (E–L) Ischemic gastrocnemius analyses at 7- and 28-days post-HLI. (E) Representative images of CD31^+^ (total) and isolectin^+^ (perfused) capillaries and quantification (F, G) in males (7dpi; control n = 6, mFATBLOCK n = 8. 28dpi; control n = 6, mFATBLOCK n = 8) and females (7dpi; control n = 8, mFATBLOCK n = 8. 28dpi; control n = 8, mFATBLOCK n = 7). (H) Representative images of Perilipin^+^ adipocytes. (I) IMAT quantification in males (7dpi; control n = 10, mFATBLOCK n = 10. 28dpi; control n = 8, mFATBLOCK n = 9) and females (7dpi; control n = 9, mFATBLOCK n = 10. 28dpi; control n = 9, mFATBLOCK n = 8). (K) Muscle wet weight in males (7dpi; n=10/genotype. 28dpi n = 8/genotype) and females (7dpi n=9/genotype and 28dpi; n=9/genotype). (L) Average myofiber cross-sectional area in males (7dpi; control n = 8, mFATBLOCK n = 7. 28dpi; n=7/genotype) and females (7dpi; control n = 8, mFATBLOCK n = 7. 28dpi; control n = 8, mFATBLOCK n = 7). Representative immunofluorescent images at 20x magnification from female mice at 7dpi, scale bars = 200 μm. Panels C and F-L were analyzed using a two-way ANOVA with Tukey’s post hoc testing for multiple comparisons when significant interactions were detected. All data are presented as mean ± SD.

Following HLI surgery, mFATBLOCK mice and littermate controls had similar paw perfusion recovery (**Figure 5D**). Seven days post-HLI, female mFATBLOCK mice had more total capillaries than their littermate controls, but this effect did not persist at day 28 (**Figure 5E, F**). This observation may suggest that female mFATBLOCK mice have more rapid ischemic angiogenic response than controls. Interestingly, this was not observed in male mice or in perfused capillary densities (**Figure 5E, G**). mFATBLOCK mice had near complete ablation of IMAT in the ischemic gastrocnemius at both 7- and 28-days post HLI (**Figure 5H, I** and **Supplemental Table 4**). Quantitatively, mFATBLOCK mice had 0.15 ± 0.10 (males) and 0.14 ± 0.07 (females) PERILIPIN^+^ area/mm^2^ of muscle area at 7-days post-HLI, whereas WT littermates had 1.60 ± 1.14 (males) and 2.13 ± 1.33 (females) PERILIPIN^+^ area/mm^2^ of muscle area. This represents a 10-15-fold reduction in IMAT levels establishing a model by which we can test whether suppressing IMAT levels can improve ischemic limb outcomes. Control mice exhibited a significant increase in FAP number following HLI, whereas mFATBLOCK mice maintained FAP levels near those of non-ischemic muscle (**Figure 5J**). Importantly, preventing FAP differentiation to adipocytes did not alter the extracellular matrix area in mFATBLOCK mice (**Supplemental Figure 7B**), indicating that fibrotic remodeling was not affected. mFATBLOCK mice had no differences in the gastrocnemius muscle weight (**Figure 5K**) but they did have significantly larger average myofiber cross-sectional areas at both timepoints compared to controls, while wet weight was not different between groups (**Figure 5L**). At 7-days post-HLI, mFATBLOCK mice had significantly fewer embryonic myosin heavy chain positive myofibers compared to littermate controls (**Supplemental Figure 8)** arguing for accelerated myogenesis. Analysis of immune cell infiltration of the ischemic gastrocnemius muscle revealed that mFATBLOCK mice had significantly fewer total leukocytes and neutrophils, but similar abundance of macrophages, when compared to littermate controls (**Supplemental Figure 9A-D**). Modest changes in whole muscle cytokine levels were also observed in mFATBLOCK mice (**Supplemental Figure 9E**).

### Ablation of IMAT improves ischemic muscle performance

Next, we tested whether preventing IMAT formation improved muscle function in mice subjected to HLI. Compared to littermate controls at both 7- and 28-days post-HLI, mFATBLOCK mice had greater muscle strength and quality evidenced by significantly higher absolute (**Figure 6A**) and specific forces (**Figure 6B**). mFATBLOCK mice also had higher muscle power output compared with littermate controls (**Figure 6C**). Because IMAT ablation resulted in a greater total area of the muscle occupied by myofibers (**Supplemental Figure 10A**), normalizing absolute forces to the total area of muscle fibers confirmed that mFATBLOCK mice had similar myofilament contractile function at 28- days post-HLI, but mFATBLOCK mice had greater myofilament contractile function at 7-days post-HLI (**Supplemental Figure 10B**). These findings are consistent with mFATGAIN mice and indicate that the early formation of IMAT following HLI delays functional recovery without causing permanent damage to the contractile apparatus. Using the 6-min limb function test, mFATBLOCK mice had significantly higher muscle work output across the test (**Figure 6D**) and the corresponding total work performed (**Figure 6E** and **Supplemental Table 4**), demonstrating that lowering IMAT formation improves ischemic muscle performance. Importantly, this improved limb function in mFATBLOCK was accompanied by significantly greater muscle perfusion during the 6-minute limb function test (**Figure 6F, G**).

**Figure 6.**
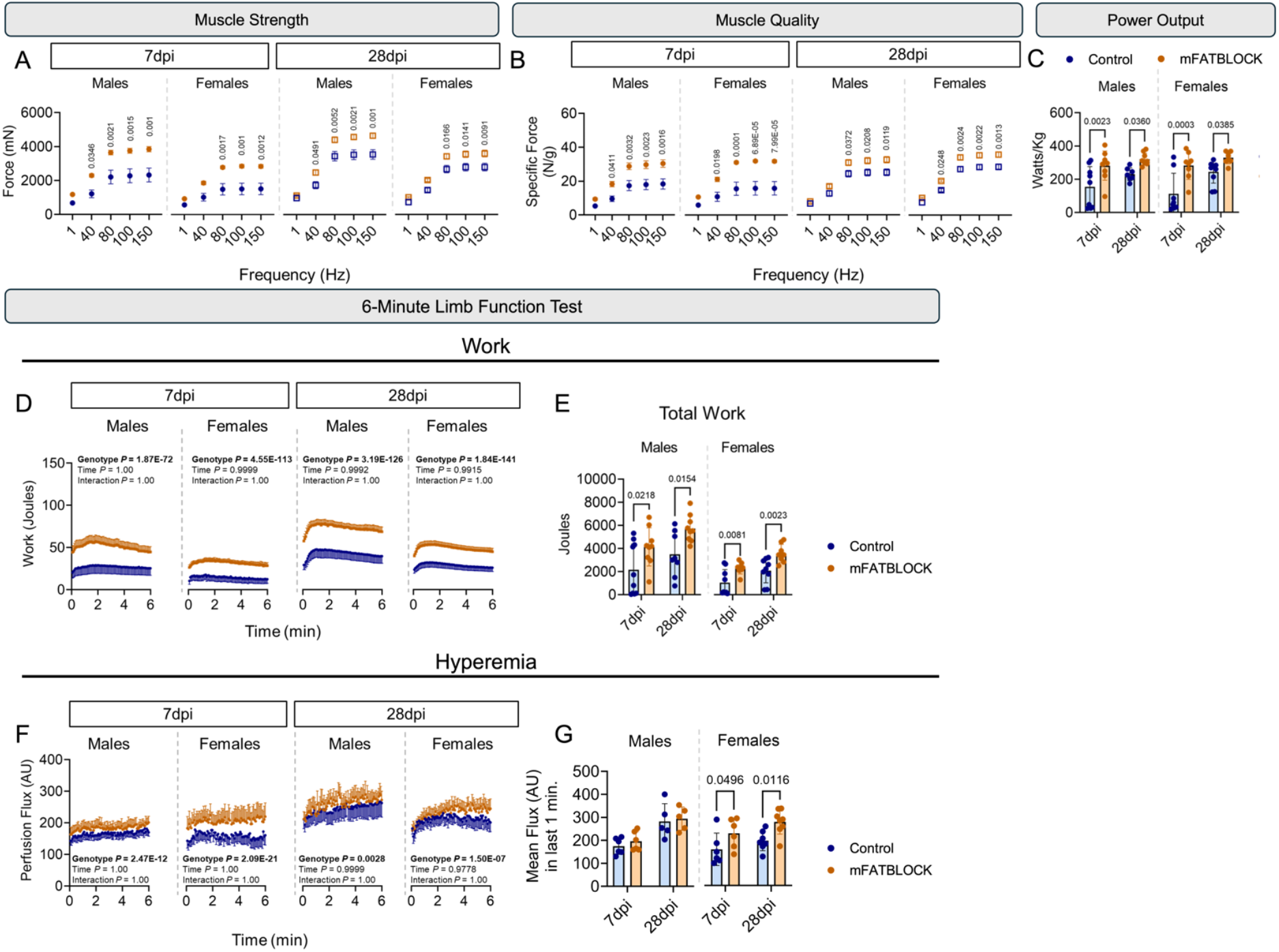
Ablation of IMAT improves function outcomes in mice with HLI. (A) Absolute Force across increasing stimulation frequencies. (B) Specific force normalized to muscle weight. (C) Power generation at 30% of 80Hz tetanic force. (D) Work performed across the 6-minute test and (E) total work performed during the test. (A-E) Males (7dpi; n=9 mice/genotype. 28dpi; control n = 8 mice, mFATBLOCK n = 9 mice) and females (7dpi; control n=9 mice mFATGAIN n=8 mice. 28dpi; control n=9 mice mFATGAIN n=8 mice). (F) Laser Doppler perfusion flux in the gastrocnemius across the test and (G) average flux during the last one minute in males (7dpi; n=6 mice/genotype. 28dpi; n=5 mice/genotype) females (7dpi; n=6 mice/genotype. 28dpi; n=8 mice/genotype). Data were analyzed using a two-way-ANOVA with Tukey’s post hoc testing for multiple comparisons and represented as mean ± SD.

### Biomarkers of IMAT accumulation correlate with ischemic muscle function in mice and humans with PAD/CLTI

Considering that genetically increasing IMAT impaired ischemic muscle function and genetically suppressing IMAT improved ischemic muscle function, we examined if the level of IMAT abundance was a primary determinant of ischemic limb function. Univariable regression analyses using the entire cohort of mice revealed that IMAT level, total myofiber area, and the mean myofiber cross-sectional area, but not hyperemia, were significantly associated with muscle strength at both 7- and 28-days post-HLI (**Figure 7A, B**). Univariable regression analyses also revealed that IMAT level, total myofiber area, and the mean myofiber cross-sectional area, were significantly associated with total work performed in the 6-minute limb function test at both 7- and 28-days post-HLI (**Figure 7C, D**). Additionally, the hyperemia achieved during the 6-minute limb function test was also significantly associated with total work performed in the 6-minute limb function test at 7-days post-HLI (**Figure 7C**). In contrast, univariable regression analyses demonstrated that absolute force and total muscular work were not significantly associated with total or perfused capillary densities, as well as paw perfusion flux at either time point (**Supplemental Figure 11**). We also performed stepwise multiple regression analysis using the data from 28-days post-HLI. Here, the model indicated that ∼58% of the variance of absolute force levels could be explained by three variables: IMAT level, average myofiber area, and total area of muscle occupied by muscle fibers (**Table 1**). Regarding total work, the stepwise multiple regression model indicated that ∼49% of the variance of total work performed could be explained by two variables: IMAT level and average myofiber area (**Table 1**). Together, these univariable and multivariable regression analyses demonstrate that IMAT levels are a key determinant of ischemic muscle function in mice with experimental PAD. Next, we aimed to explore if a similar relationship may exist in patients with and without PAD/CLTI. Using data from a recently published cohort^30^, we examined the association between plantarflexor muscle strength and adipogenic gene expression (*FABP4, CEBPA*, and *ADIPOQ*) as a surrogate for IMAT abundance since the muscle biopsies for this cohort were not fixed and stored in a manner to allow adipocyte quantification. Nonetheless, the expression of *FABP4, CEBPA*, and *ADIPOQ* all had statistically significant negative correlation with plantarflexor strength in humans with and without PAD/CLTI (**Figure 7E**). Plantarflexor strength was also strongly correlated with ankle-brachial index (Pearson *r*=0.65, *P*=1.83E-07), indicating that more severe PAD/CLTI is associated with weaker calf muscle strength. Adipogenic gene expression (*FABP4, CEBPA*, and *ADIPOQ*) also had a statistically significant negative correlation with ankle-brachial index, suggesting that more severe limb ischemia promotes greater IMAT abundance (**Figure 7F**). Together, these data suggest that individuals with more IMAT formation will have lower muscle function and that strategies which can lower IMAT levels may hold potential to improve limb function and possibly walking performance in patients with PAD/CLTI.

**Figure 7.**
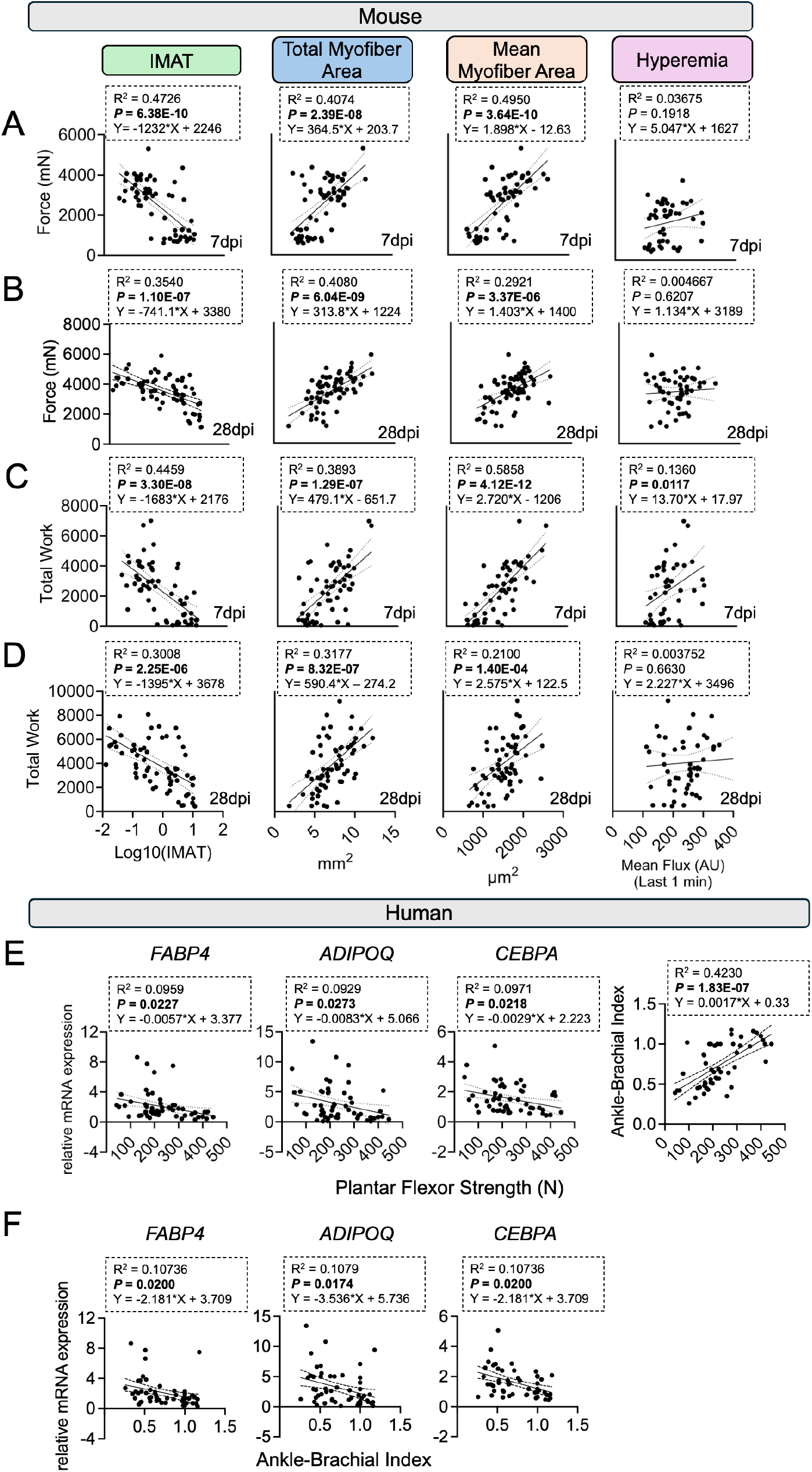
Increased adipogenesis is correlated with limb functional outcomes in human and murine ischemic limbs: Limb function data were pooled data from all mFATGAIN, mFATBLOCK, and corresponding controls (A) Univariable regression analysis for absolute force levels 7-days post-HLI. (B) Univariable regression analysis for total work performed 7-days post-HLI. (C) Univariable regression analysis for absolute force levels 28-days post-HLI. (D) Univariable regression analysis for total work performed 28-days post-HLI. (E-F) Regression analysis between plantarflexor strength, adipogenic gene expression levels, and ankle-brachial index in patients with PAD/CLTI and non-PAD control participants (n=54). All data were analyzed using a simple linear regression and Pearson correlation.

**Table 1:** Multivariate regression analyses for ischemic muscle function.

| **Model – Absolute Force at 28-days post HLI** | **Adjusted $\beta$** | ***P*-value** | **Partial $R^2$** | **VIF** | **Model $R^2$** | **Adjusted $R^2$** |
| --- | --- | --- | --- | --- | --- | --- |
| (Constant) |  | 0.020 |  |  | 0.608 | 0.583 |
| IMAT | -0.421 | 7.82E-04 | 0.462 | 1.66 |  |  |
| Average myofiber area | 0.312 | 0.002 | 0.430 | 1.09 |  |  |
| Total myofiber area | 0.272 | 0.025 | 0.319 | 1.66 |  |  |
| **Model – Total Work at 28-days post HLI** | **Adjusted $\beta$** | ***P*-value** | **Partial $R^2$** | **VIF** | **Model $R^2$** | **Adjusted $R^2$** |
| (Constant) |  | 0.286 |  |  | 0.509 | 0.488 |
| IMAT | -0.588 | 2.96E-06 | 0.608 | 1.08 |  |  |
| Average myofiber area | 0.318 | 0.004 | 0.400 | 1.08 |  |  |

## DISCUSSION

The studies herein demonstrate that patients with PAD/CLTI present with a marked accumulation of IMAT and increased adipogenic signatures within FAPs in muscle specimens collected from the diseased limb. Notably, this phenotype is remarkably different from patients with mild PAD (intermittent claudication) or non-PAD participants, implicating IMAT as an underlying mechanism contributing to the limb pathobiology in PAD/CLTI. Importantly, the IMAT phenotype of PAD/CLTI limb muscle is present across numerous independent patient datasets, collected by independent labs, and detected using varied quantitative approaches including qPCR, RNA sequencing, single cell/nucleus RNA sequencing, proteomics, and lipidomics^28,30,43,44,47^, demonstrating the fidelity of this finding. Using rigorous genetic gain- and loss-of-function models to both increase and decrease IMAT levels, we demonstrate that the level of IMAT present directly impacts muscle strength and performance in the ischemic limb.

Because the root cause of PAD/CLTI originates from atherosclerotic occlusion and/or narrowing of the arteries, patients with the disease are at higher risk for major cardiovascular events and limb amputation. As such, medical guidelines currently focus on reducing these risks through lifestyle interventions (exercise) and usage of antithrombotic, anticoagulant, and antiplatelet therapies. Less progress has been made at improving lower limb function and walking performance. In fact, a recent statement by the Society for Vascular Medicine outlined developing novel approaches to improve walking performance as a key research priority for PAD^51^. Using conditional deletion of Pparγ in FAPs, we demonstrate that repressing IMAT formation in mice improves the total work performed in a 6-min limb function test, an outcome akin to the distance walked by patients during a 6-min test. On the other hand, conditional overexpression of Pparγ in FAPs significantly reduced the total work performed in a 6-min limb function test. These findings indicate that developing clinical strategies to limit or reverse IMAT formation in the limb muscle of patients with PAD/CLTI should improve the limb muscle function – addressing a critical need for patients. Our findings point to Pparγ as an exciting target to modulate IMAT in PAD/CLTI. However, therapeutic targeting of Pparγ is not a trivial approach. Our data would suggest that Pparγ antagonists might improve limb function in PAD/CLTI. However, systemic Pparγ inhibition could have unwanted systemic metabolic side effects, like those seen in patients with pathological variants in Pparγ causing lipodystrophy^52^. On the other hand, some studies have reported that Pparγ agonists (such as pioglitazone) or pan-Ppar agonists (such as bezafibrates), improve ischemic angiogenesis/neovascularization^53-55^, although it has been reported that these vascular effects occur independent of Pparγ^56^. While it is unknown whether pioglitazone had any impact on IMAT in those studies, based on our genetic gain- and loss-of-function models, we would hypothesize that pharmacological activation of Pparγ would increase IMAT formation and worsen ischemic limb function. In addition, Pparγ is required by multiple cell types in the ischemic muscle, including endothelial cells and immune cells; thus, the effects of systemic Pparγ modulation are likely pleiotropic, simultaneously beneficial in some cells and detrimental in others. To this end, treatment of mice with GW9662, a Pparγ antagonist failed to repress IMAT formation following a non-ischemic glycerol injection muscle injury (**Supplemental Figure 15**). Thus, while targeting IMAT represents an exciting new therapeutic strategy for PAD/CLTI, future work is required to determine whether manipulating Pparγ itself is the right approach, or whether more selective anti-adipogenic pathways can be identified.

Previous studies have confirmed that FAPs are the primary source of ectopic adipocyte formation in skeletal muscle following acute non-ischemic injuries ^22,57-59^. However, it is important to note that these mesenchymal stromal cells also facilitate myogenesis^60^ and depletion of FAPs impairs muscle regeneration^61^. Thus, an important finding from the current study was that FAP abundance was unchanged in both mFATGAIN and mFATBLOCK mice and under non-ischemic conditions these manipulations of Pparγ in FAPs did not alter the level of IMAT, myofiber size, or capillary densities. We report that PPARγ overexpression in FAPs is not sufficient to force adipogenic differentiation of FAPs without ischemic injury, indicating that there are additional molecular ‘brakes’ that prevent adipogenic differentiation in non-injured conditions. Another important aspect of this study is that neither model displayed systemic change in body weight, body fat percentage, or glucose homeostasis thereby avoiding systemic effects that occur in other fat ablation models^62,63^.

Our experiments reveal that adipocytes form rapidly after the onset of limb ischemia and are visible as early as two days post-HLI. In both mouse experiments, we observed no change in the IMAT area between 7-, 28-, and 70-days post-HLI demonstrating that once IMAT forms in the ischemic limb, it remains present despite recovery of limb perfusion (**Supplemental Figures 12 and 13**). Temporal analyses at 7- and 28-days post-HLI allow some interpretation regarding how IMAT formation impacts the functional recovery of muscle following an ischemic injury. Mice with more IMAT exhibited lower muscle strength. When normalizing muscle forces to the total muscle fiber area, we observed relatively normal specific forces at 28-days post-HLI (**Supplemental Figures 5 and 10**). This finding suggests that the contractile proteins present in the limb are functioning normally (i.e., intrinsic function of myofilament proteins in normal), but the early formation of IMAT reduces the number of contractile elements within the ischemic muscle. However, at 7-days post HLI, having more IMAT impaired the intrinsic contractile function (**Supplemental Figures 5 and 10**) suggesting that the early formation of adipocytes delays the functional regeneration of ischemic muscle. Consistent with this interpretation, the amount of IMAT at 7-days post-HLI was associated with the amount of myofibers expressing embryonic rather than mature isoforms of the myofilament protein - myosin heavy chain (**Supplemental Figures 3 and 8**). These findings are congruent with our observations made using non-ischemic muscle injury models where IMAT formation restricted the early muscle regeneration culminating in impaired muscle function^36^.

Historically, most preclinical HLI studies have used laser Doppler perfusion recovery as the primary outcome measure due to being non-invasive and highly repeatable. Most groups focus laser Doppler perfusion recovery measures on the footpad/paw because this area of the limb is hairless and the prone positioning of the mouse results in high reproducibility^64^. Additionally, many assume that the most distal tissues like the paw experience the most severe ischemic condition following HLI. However, recent findings have questioned whether this is the best approach for preclinical HLI studies. For example, Amorese *et. al*. ^65^ demonstrated that the flexor digitorum brevis (FDB) muscle is uniquely protected from ischemia compared to other hindlimb muscles. Here the authors linked the hypoxia resistance of the FDB muscle to unique metabolic adaptations that included overexpression of GLUT1, however GLUT1 overexpression was not sufficient to induce hypoxia resistance in other hindlimb muscles. Unexpectedly, we found that the FDB muscle did not form IMAT following hindlimb ischemia, including the mFATGAIN mice (**Supplemental Figure 14A, B**). This striking phenotype occurred despite the FDB muscle having a greater relative abundance of FAPs compared to the gastrocnemius muscle (**Supplemental Figure 14C, D**). The mechanisms underlying the lack of IMAT formation in the FDB muscle are not fully known but could be related to the enhanced ability of the muscle to survive the ischemic insult with HLI. This concept is supported by data demonstrating that when muscle regeneration is genetically repressed, IMAT formation is amplified following HLI^66^. Regardless, this observation adds the growing evidence that paw perfusion recovery may not be the best outcome for HLI studies and reinforces the strengths of using the 6-min limb function test^40^ to assess limb function in these models. To this end, univariable and multivariable regression analyses also demonstrated that the level of IMAT was a key determinant of muscle strength and total work performed, whereas resting paw perfusion failed to associate with these limb function outcomes.

We acknowledge several limitations of the current studies. First, the HLI model is an acute, surgically induced ischemic event, which does not replicate the chronic and progressive nature of the atherosclerotic occlusion seen human patients with PAD/CLTI. However, HLI models result in a highly reproducible ischemic injury that can be useful in studying the pathobiology of the ischemic tissues ^67^. Additionally, experiments herein used young mice lacking comorbidities such as hyperlipidemia or diabetes, which are often found in patients with PAD/CLTI. However, previous studies suggest that the inclusion of these risk factors is likely to enhance IMAT formation^68-70^. Second, human specimens used to correlate adipogenic gene expression signatures with plantarflexor strength were not processed in a manner to allow accurate quantification of IMAT. Because of human tissue processing procedures, we also did not assess the degree of muscle fibrosis, atrophy, or inflammation which may influence gene expression and muscle function outcomes thereby influencing the correlation strength. Moreover, while the ankle-brachial index was negatively correlated with adipogenic gene expression in humans, this is not the most sensitive hemodynamic measure and has considerable variance across patients. Additionally, it does not reflect the microvascular perfusion to the muscle. Thus, these significantly correlations should be interpreted with caution and future studies should quantify IMAT and hemodynamics with greater precision. Nonetheless, prior cohort studies employing CT imaging approaches have linked IMAT accumulation to leg function in patients with PAD^41^, a finding that supports the correlation results we observed. Beyond the context of CLTI, the formation of IMAT and its role in determining limb function may also be relevant to acute limb ischemia where our findings herein would suggest that the acute ischemic event is likely to provoke FAP-to-adipocyte differentiation like the observed outcomes in mice with HLI.

While the current study demonstrates that IMAT formation occurs with ischemic injury and that the level of IMAT formation is a key determinant is ischemic limb function, several important questions remain unanswered. First, while the cellular source of IMAT is predominantly the FAPs, the initiating signal for adipogenic differentiation is not clear, particularly in the complex microenvironment of the PAD/CLTI limb. Hedgehog signaling has been reported to control FAP-adipocyte differentiation^58,71^, however, whether this pathway is altered in human patients with PAD/CLTI has not been reported. Previous studies have shown that modulation of the Hedgehog pathway can promote muscle recovery and angiogenesis in mice with hindlimb ischemia ^72-77^ suggesting that targeting this pathway could have pleiotropic benefits in PAD/CLTI. Recent work using single cell and single nucleus RNA sequencing showed that FAPs and immune cells are highly communicative in the muscle of patients with PAD^33,78^. We observed that when IMAT levels were high ischemic muscles had greater abundance of total leukocytes, macrophages, and neutrophils. In contrast, blocking IMAT formation significantly lowered total leukocyte and neutrophil abundance, but did not change the number of macrophages in the ischemic muscle. These observations could indicate that immune cell-FAP intercellular communication plays a role in IMAT formation. Arguments against this mechanism include the observation that adipocytes are found as early as 2-days post-HLI (Figure 1) which is before significant immune cell infiltration occurs, and that immunodeficient mice subjected to hindlimb ischemia display visible adipocytes in histological analyses^79^. On the other hand, the changes in immune cell abundance with IMAT manipulation could indicate increased immune cell abundance is a reactionary event caused by increased IMAT formation. The current study cannot rule out the alternative hypothesis in which immune cell-FAP interactions may causally drive adipocyte differentiation and IMAT accumulation in the ischemic limb muscle. Clearly, additional work is needed to fully elucidate the triggering mechanism(s) that promote FAP differentiation into IMAT with ischemic injury and determine whether immune cells play a causal role in IMAT accumulation. Second, while the mFATBLOCK model is a rigorous genetic approach, this model prevented the formation of IMAT. More studies are needed to identify mechanisms to reverse or shrink IMAT levels as these approaches represent a more viable therapeutic approach for patients with PAD/CLTI who are often seen clinically long after the onset of the first ischemic events. Again, caution is needed in developing these approaches because FAPs are important for muscle homeostasis so treatment approaches should repress the adipogenic potential without impacting pro-myogenic functions of FAPs.

In conclusion, this study establishes that IMAT accumulation is a hallmark of muscle pathology in patients with PAD/CLTI. Using rigorous genetic models, we demonstrate a direct link between FAP-derived IMAT accumulation and ischemic limb function in mice with experimental PAD/CLTI. Together, these studies demonstrate that IMAT is not merely a histological feature of disease but a functional contributor of poor limb function and provide evidence that strategies capable of limiting or reducing IMAT accumulation hold promise for improving limb function in PAD/CLTI.

## ABBREVIATIONS

CLTI: chronic limb threatening ischemia
DEG: differentially expressed gene(s)
FAPs: fibro-adipogenic progenitor cells
GSEA: gene set enrichment analysis
IMAT: intramuscular adipose tissue
PAD: peripheral arterial disease
ROS: reactive oxygen species
snRNAseq: single nucleus RNA sequencing

## Acknowledgements

Raw supporting data for this paper can be found at: 10.6084/m9.figshare.30521537.

