## Supplemental Material for "Intramuscular Adipose Tissue Accumulation is a Key Determinant of Limb Function in Peripheral Artery Disease"

[2]Department of Surgery

[4]Myology Institute

[7]Department of Physiology

Short Title: IMAT regulates muscle function in PAD

[#]Correspondence:

Terence E. Ryan, PhD:

Daniel Kopinke, PhD:

**KEYWORDS:** limb ischemia, muscle, fibro-adipogenic progenitor, FAPs, stem cell

**Detailed Materials and Methods**:

Human Studies and Datasets**.** Gastrocnemius muscle specimens were collected from PAD/CLTI and non-PAD patients via percutaneous muscle biopsy using sterile procedures as previously described [1,2] and used to generate total RNA for gene expression analysis. This study was approved by the institutional review boards at the University of Florida and the Malcom Randall VA Medical Center (Protocol IRB201801553). All study procedures were carried out according to the Declaration of Helsinki and participants were fully informed about the research and informed consent was obtained. A detailed description of the patient cohort has been previously published[3]. We also analyzed several publicly available and previously published datasets involving human patients with PAD/CLTI. Bulk RNA sequencing data from the gastrocnemius muscle of non-PAD, patients with intermittent claudication, and patients with CLTI undergoing limb amputation was obtained here[1] (GSE120642). Proteomic data from non-PAD volunteers and patients with CLTI was obtained here[2] (PXD021849). Lipidomic data from non-PAD volunteers and patients with CLTI was obtained here[4] (Metabolomics Workbench Study ID's: ST001615, ST001616, ST001617). Single-cell RNA sequencing data from patients with CLTI was obtained from Southerland *et al.*[5] (GSE227077). Single-nucleus RNA sequencing data from non-PAD volunteers and patients with PAD/CLTI was obtained here[3,6] (GSE233882 and GSE284482).

Analysis of Single Cell and Single Nucleus RNA sequencing Datasets. We used single cell and single nucleus RNA sequencing datasets that have been previously published[3,5,6]. Each of the publications describe the detailed processing and annotation of the datasets. In this paper, we subset the fibro-adipogenic progenitor cells (FAPs) in each dataset using Scanpy. Target gene expression comparisons within the FAPs were made using a Wilcoxon test.

Animal Studies. All animal experiments adhered to the *Guide for the Care and Use of Laboratory Animals* from the Institute for Laboratory Animal Research, National Research Council, Washington, D.C., National Academy Press. All procedures were approved by the Institutional Animal Care and Use Committee of the University of Florida (Protocol 202300000490). Mice were housed in a temperature- (22°C) and light-controlled (12:12-h light-dark) room and maintained on standard chow (Envigo Teklad Global 18% Protein Rodent Diet 2918 irradiated pellet) with free access to food and water. BALB/cJ mice (Stock No. 000651) were obtained from Jackson Laboratories. To manipulate IMAT formation, we targeted the transcription factor *Pparγ,* a key regulator of adipogenesis, using a tamoxifen-inducible, fibro-adipogenic progenitor cell (FAP) specific *Pdgfrα$^{CreERT2}$* mouse (Jackson Laboratories, Stock No. 032770)[7]. *Pdgfrα$^{CreERT2}$* mice were bred with conditional transgenic mice expressing a human wild-type PPARG allele under control of the CAG promoter (CAG-LSL-PPARG; kindly provided by Dr. Curt Sigmund, Medical College of Wisconsin)[8] to generate a FAP-specific, inducible overexpression of PPARγ model termed mFATGAIN. To block IMAT formation, crossed the *Pdgfrα$^{CreERT2}$* mice with floxed *Pparg* mice (B6.129-*Ppargtm2Rev*/J (PPARγ$^{loxP}$, Jackson Laboratories Stock No. 004584) carrying an inducible EYFP reporter (B6.129X1-*Gt(ROSA)26Sortm1(EYFP)Cos*/J, Jackson Laboratories Stock No. 006148) which we termed mFATBLOCK[9]. All FAP-specific mouse lines were backcrossed a minimum of three generations to the 129S1/SvlmJ genetic background because C57BL/6J mice are resistant to limb ischemia and form little IMAT with non-ischemic injuries[10]. In all experiments, littermates lacking the Cre transgene were used as controls. All experiments involved male and female mice that were randomized to experimental groups so that both the surgeon and experimenters were blinded to the treatment groups and genotypes. Female mice underwent an ovariectomy two weeks prior to enrollment. All mice received tamoxifen (200-250 mg/kg, Cat No. T5648) dissolved in sunflower seed oil via oral gavage for three consecutive days to induce Cre recombination. Two weeks after tamoxifen administration, femoral artery ligation was performed to induce unilateral hindlimb ischemia (HLI)[11,12].

Blood Glucose Analysis. Blood glucose was analyzed in fed, conscious mice by collecting a small blood sample from a tail snip directly on a glucose test strip. All measurements were performed between 8:00 – 9:00am.

Body Composition. Body composition was determined using EchoMRI and dual-energy X-ray absorptiometry (InAnalyzer 2, Micro Photonics, Inc.) according to the manufacturer's instructions.

Primary FAP isolation. FAPs were isolated by differential plating as previously described [9,13] with some modifications. In brief, hindlimb muscles were carefully dissected and mechanically dissociated. Muscle was then digested using 0.15% w/v Collagenase type II (Gibco, Cat No. 17101-015), 0.04% Dispase II (Sigma, Cat No. D4693) in Dulbecco's Modified Eagle Medium (DMEM) for 30min at 37C. A solution containing 2mM Ethylenediaminetetraacetic acid (EDTA) and 0.5% bovine serum albumin (BSA) in phosphate buffered saline (PBS) was added to stop enzymatic digestion and the suspension was sequentially filtered through 100 µm and 40 µm strainers. Cells were centrifuged at 800 g for 10 minutes at 4 °C, resuspended in DMEM, and plated in a T-75 flask. After allowing cells to adhere for one hour at 37 °C, non-adherent cells were discarded and adherent cells (FAPs) were collected for analysis.

RNA Isolation and Quantitative Polymerase Chain Reaction (PCR). All samples were lysed with TRIzol reagent (Invitrogen, Cat. No. 15-596-018). Muscle biopsies were homogenized using a PowerLyzer 24 bead mill homogenizer (Qiagen). Total RNA was purified with the Direct-zol RNA MiniPrep Kit (Zymo Research, Cat. No. R2052) according to the manufacturer's instructions. LunaScript RT SuperMix Kit (New England Biolabs, Cat. No. E3010L) was used for cDNA generation. Quantitative PCR was performed using the Luna Universal qPCR Master Mix (New England Biolabs,

Cat. No. M3003X) with SYBR Green-based detection on a QuantStudio 3 Real-Time PCR System (ThermoFisher Scientific). Relative gene expression was calculated using the $2^{-\Delta\Delta CT}$ method with expression normalized to the respective control group of each experiment. Primers are used are available in Supplemental Table 1.

Hindlimb Ischemia. Femoral artery ligation was performed by placing silk ligatures on the femoral artery just distal the inguinal ligament and immediately proximal to the saphenous and popliteal branches to surgically induce unilateral hindlimb ischemia (HLI) [11,12]. Mice were anesthetized with intraperitoneal injection of ketamine (100 mg/kg) and xylazine (10 mg/kg). Mice were given a subcutaneous injection of extended-release buprenorphine (Ethiqa, 3.25 mg/kg) prior to surgery for analgesia. Given that most female patients with PAD are post-menopausal, female mice underwent a bilateral ovariectomy 7-10 days prior to study enrollment. For ovariectomy, mice received subcutaneous injections of extended-release buprenorphine (3.25 mg/kg, prior to surgery) and meloxicam (15-20 mg/kg, prior to surgery and once every 24 hours for 72 hours post-operatively) for analgesia.

Laser Doppler Perfusion Imaging. Perfusion recovery was measured using laser Doppler perfusion imaging (LDPI) (Model LDI2-IR, Moor Instruments, Wilmington, DE). Mice were anesthetized with a mixture of ketamine (100 mg/kg) and xylazine (10 mg/kg) and placed under the scanner in a prone position on a temperature-controlled heating pad. Data were analyzed using MoorLDI Review Software (V6.2) as previously described[14] and perfusion was reported as a percentage of the non-ischemic limb.

Labeling of Perfused Capillaries. To visualize perfused capillaries, mice received a retro-orbital injection of 50 µL *Griffonia simplicifolia* isolectin B4-DyLight 649 (1 mg/mL, Vector Laboratories, Cat No. DL-1208). Following injection, animals were returned to their cages and allowed 1–2 hours of free movement to permit systemic circulation of the labeled isolectin prior to terminal experiments.

Skeletal muscle histology. Human skeletal muscle biopsies were embedded in optimal cutting temperature (OCT) compound, frozen in liquid nitrogen–cooled isopentane, and cryosectioned at 5 µm using a Leica cryostat (Model No. 3050S) and mounted onto microscope slides. Slides were incubated in Mayer's hematoxylin solution (Millipore-Sigma, Cat. No. MHS1) for 15-minutes followed by a 15-minute wash in circulating, lukewarm water. Slides were then submerged in eosin (Millipore-Sigma, Cat. No. HT110132) for one minute then immediately dipped five times in increasing concentrations of ethanol (60% - 100%). Slides were cleared using a 1:1 xylene:ethanol solution and cover slipped with Permount mounting media (ThermoFisher Scientific, Cat. No. SP15-100).

Muscles from mice were fixed in 4% paraformaldehyde (ThermoFisher Scientific, Cat. No. J61899.AK) for 3–4 hours at 4°C, rinsed in PBS, and cryoprotected overnight in 30% sucrose. Tissues were embedded in OCT compound, frozen in liquid nitrogen–cooled isopentane, and cryosectioned at 10 µm using a Leica cryostat (Model No. 3050S). For all immunofluorescence staining, sections were blocked and permeabilized using a solution of 5% goat serum, 0.3% Triton X-100, and 1% BSA in 1× PBS. For quantification of total capillaries, sections were stained with rabbit anti-CD31 primary antibody (1:100, Abcam, Cat. No. ab28364) and Alexa Fluor 555–conjugated goat anti-rabbit secondary antibody (1:250, ThermoFisher Scientific, Cat. No. A21428). FAPs were labeled with goat anti-PDGFRα primary antibody (1:250, R&D Systems, Cat. No. AF1062) and Alexa Fluor 555–conjugated donkey anti-goat (1:1000, ThermoFisher Scientific, Cat. No. A21432), along with chicken anti-GFP (1:1000, Aves Labs, Cat. No. GFP-1020) to visualize the EYFP endogenous reporter. Intramuscular adipose tissue (IMAT) was detected using rabbit anti-Perilipin-1 primary antibody (1:1000, Cell Signaling, Cat. No. 9349S) and Alexa Fluor 488–conjugated goat anti-rabbit secondary antibody (1:250, ThermoFisher Scientific, Cat. No. A11008), on the same slide, myofibers were visualized using Phalloidin conjugated with Alexa Fluor 594 (1:1000, ThermoFisher Scientific, Cat. No. A12381). Wheat germ agglutin (WGA) conjugated with Alexa-Flour488 (1:100, ThermoFisher Scientific, Cat. No. W11261) was used to label the extracellular matrix. To quantify regenerating myofibers, muscle sections were labeled with a primary antibody for embryonic myosin heavy chain (MYH3; Developmental Studies Hybridoma Bank, University of Iowa, Cat. No. F1.652, 1:100 dilution) overnight at 4C. Following washes, the sections were then labeled with Alexa Fluor 555–conjugated goat anti-mouse (1:1000, ThermoFisher Scientific Cat. No. A21422) secondary antibody. The number of MYH3$^{+}$ myofibers were quantified by manually counting in ImageJ. Immune cells were detected using rat-anti-CD45 (1:500, eBioscience, Cat. No. 14045182), rat anti-Ly6G (1:100, ThermoFisher Scientific, Cat. No. A21422), and mouse anti-CD68 (1:100, ThermoFisher Scientific, Cat. No. 501122651) primary antibodies. Following washes, the cells were visualized by labeling with Alexa Fluor 555–conjugated goat anti-rat (1:1000, ThermoFisher Scientific Cat. No. A21434), and Alexa Fluor 555–conjugated goat anti-mouse (1:1000, ThermoFisher Scientific Cat. No. A21422) to label leukocytes, neutrophils, and macrophages respectively. The abundance of immune cells were quantified by manually counting in ImageJ.  All primary antibodies were applied overnight at 4°C. The following day, slides were washed in 1× PBS and all secondary antibodies or directly conjugated dyes were applied for 1.5 hours at room temperature. Coverslips were mounted onto slides using Vectashield hardmount with DAPI (Vector Laboratories, Cat. No. H-1500). All slides were imaged using an Evos FL2 Auto microscope (ThermoFisher Scientific) at 20x magnification, and tiled images of the entire muscles were obtained for analysis.

Histological Analysis. All images were processed the same for each sample by a blinded investigator. FAPs were quantified by manually counting PDGFRα$^{+}$ cells in 2-3 randomly selected 20x images from each head of the gastrocnemius (5-6 total images per animal) and in stitched 20x images of the entire flexor digitorum brevis muscle. The number of FAPs was normalized to the total muscle area. Myofiber cross-sectional area was quantified from phalloidin-stained images using a previously described pipeline in which images are segmented with Cellpose and area is measured using the ImageJ plugin LabelsToRoi [15]. Individual fiber areas within a cryosection were summed to calculate total area of the muscle occupied by myofibers. IMAT was quantified by thresholding Perilipin-stained sections to measure the total Perilipin$^{+}$ area, which was normalized to the total muscle area. For capillary and leukocyte quantifications, total (CD31$^{+}$) and perfused (Isolectin$^{+}$) capillaries, or total leukocytes (CD45$^{+}$) were calculated from thresholded images using the particle analysis tool in Fiji/ImageJ and normalized to total muscle area. Perfused capillaries are reported as a percentage of total capillaries. Fluorescence images stained with WGA and perfused isolectin, were used to quantify extracellular matrix area while excluding capillary signal. Individual channels were thresholded separately to generate binary masks. The Isolectin$^{+}$ area was subtracted from the WGA mask using the Image Calculator function in Fiji/ImageJ, and the resulting area was measured and normalized to total image area. Neutrophils and macrophages were counted manually in 2-3 randomly selected 20x images from each head of the gastrocnemius (5-6 total images per animal) and normalized to total analyzed area. Regenerating myofibers are reported as the proportion of eMyHC expressing fibers using the entire muscle section.

Cytokine Analysis. Snap frozen soleus and plantaris muscles were homogenized in 300-400ul of RIPA buffer (ThermoFisher Scientific, Cat. No. 89901) using a bead beater (PowerLyzer 24 Homogenizer, Qiagen, 110/220V). The muscle homogenate was centrifuged at 1000 x g for 10 minutes at 4°C and the supernatant was collected for cytokine analysis. We used Luminex® xMAP® technology to quantitatively and simultaneously detect fourteen mouse cytokines, chemokines and growth factors were measured. The multiplexing analysis was performed by Eve Technologies Corporation (Calgary, Alberta, Canada) using the Luminex® 200™ system (Luminex Corporation/DiaSorin, Saluggia, Italy) with Bio-Plex Manager™ software (Bio-Rad Laboratories Inc.,

Hercules, California, USA). Fourteen markers were measured in the samples using the Eve Technologies' Mouse Cytokine/Chemokine Focused 14-Plex Discovery Assay® Array (MDF14) as per the manufacturer's instructions for use (MILLIPLEX® Mouse Cytokine/Chemokine Magnetic Bead Panel Cat. #MCYT1-190K, MilliporeSigma, Burlington, Massachusetts, USA). The 14-plex consisted of GM-CSF, IFNγ, IL-1β, IL-2, IL-4, IL-5, IL-6, IL-10, IL-12p70, IL-13, IL-17A, IL-17F, MCP-1/CCL2, and TNFα. Assay sensitivities of these markers range from 0.52 – 6.61 pg/mL. Individual analyte sensitivity values are available in the MilliporeSigma MILLIPLEX® protocol.

Muscle Function Analysis. *In situ* muscle function was assessed in the plantar flexor complex (Aurora Scientific, model 1300A), as previously described [16]. Mice were anesthetized (ketamine 100 mg/kg, xylazine 10 mg/kg) and the gastrocnemius/soleus/plantaris complex of the ischemic limb was carefully isolated from its distal insertion without disrupting the vasculature. The distal *Achilles* tendon was secured with a 4-0 silk suture and attached to the lever arm of a force transducer (Cambridge Technology, Model 6650LR). The sciatic nerve was stimulated through bipolar electrodes using square-wave pulses (Aurora Scientific, Model 701A stimulator), and data were collected using a LabVIEW-based DMC system (version 615A.v6.0, Aurora Scientific Inc.). All functional tests were conducted at the optimal muscle length determined individually for each mouse using twitch contractions. Isometric contractions were elicited with 500-ms trains (2 mA, 0.2-ms pulse width) to conduct a force-frequency protocol at stimulation frequencies of 1, 40, 80, and 150Hz, with 1-minute rest intervals between contractions. Maximum tetanic force at each frequency was reported as absolute force, specific force (normalized to muscle mass), and myofiber-normalized force (normalized to total myofiber area). The peak tetanic force generated at 80Hz was recorded and used as the reference force for the 6-minute limb function test as described below.

Following a 3-minute recovery period, we performed a 6-minute limb function test to comprehensively assess muscle function. Detailed methods of the test are described in a previous publication[16]. Briefly, the plantar flexor complex was stimulated at 80Hz and the lever arm was allowed to shorten once the force exceed 30% of the absolute force level produced during the 80Hz isometric contraction. This isotonic (shortening) contraction was followed by a 2.5-second static rest period during which perfusion was quantified in the gastrocnemius muscle using laser Doppler flowmetry (moorVMS-LDF, Moor Instruments). Immediately following the static perfusion measurement, the muscle underwent three small passive stretches as previously described. This protocol produces a contraction every ~4 seconds which was repeated for the entire 6-minute test period. The average perfusion flux for each motionless rest period was quantified and plotted over time. Power, shortening velocity, and mechanical work were calculated for every contraction and plotted across time. The total work performed was calculated by summing the work of each individual contraction. Data from the 6-minute limb function test were analyzed using custom scripts in MATLAB (MathWorks, Natick, MA).

<u>Glycerol muscle injury and Pparg antagonism experiment.</u> Non-ischemic muscle injuries were performed on 8–12-week-old 129Sv/J mice (4 males in control, 5 males in treatment group) under anesthesia with isoflurane where the tibialis anterior muscle was injected with 30-50 µL of 50% glycerol (GLY; Acros Organics, 56-81-5) diluted in sterile saline. Following the glycerol injury mice were treated once daily on days 1-4 with intraperitoneal injections of the Pparγ antagonist GW9662 (1mg/kg; MedChemExpress, Cat. No.: HY-16578). GW9662 was first dissolved in dimethyl sulfoxide (DMSO) (100 mg/ml) and diluted 1:500 in sterile saline for injections. Control mice were treated with equal volumes of the DMSO/saline vehicle. The tibialis anterior muscle of both legs were harvested at day 7 post-glycerol injection for immunohistology analysis to quantify IMAT and myofiber areas.

<u>Statistical Analysis.</u> Normality of data was tested with the Shapiro-Wilk test and inspection of QQ plots. Data are presented as mean ± standard error of the mean (SEM) unless otherwise indicated.

For comparisons between two groups, Student's t-tests were used when data were normally distributed. If normality could not be established, or sample size was < 6, the non-parametric Mann-Whitney test was applied. Two-way ANOVA was used to analyze the effects of multiple factors across experimental groups with Tukey's post hoc testing for multiple comparisons when significant interactions were detected. Perfusion recovery over time was analyzed using mixed-effects models to account for repeated measures. Simple linear regression and Pearson correlation were used to assess individual relationships between myofiber/vascular properties and functional outcomes. All statistical testing above was conducted using GraphPad Prism software (version 10.0). To determine which measures were associated with the greatest functional improvements (total work), we performed stepwise multiple linear regression analyses using SPSS (version 29.0.2.0). Normality of the residual distribution was assessed by inspection of histograms. For all analyses, $P < 0.05$ was considered statistically significant.

**Supplemental Table 1: Primers used in the study.**

| qPCR Primers | | | |
|---|---|---|---|
| | | Sequence (5' -> 3') | |
| **Species** | **Gene ID** | **Forward** | **Reverse** |
| Homo Sapiens | *PPARG* | GCTTGGGTCGGCCTCG | TGGCTTCTTTCAAATCTGCGG |
| Mus Musculus | *Fabp4* | AAGGTGAAGAGCATCATAACCCT | TCACGCCTTTCATAACACATTCC |
| Mus Musculus | *Pparg* | TCTTCCATCACGGAGAGGTC | GATGCACTGCCTATGAGCAC |
| Mus Musculus | *Cebpa* | CAAGAACAGCAACGAGTACCG | GTCACTGGTCAACTCCAGCAC |
| Homo Sapiens | *ADIPOQ* | GCAGTCTGTGGTTCTGATTCCATAC | GCCCTTGAGTCGTGGTTTCC |
| Homo Sapiens | *FABP4* | ATGGGATGGAAAATCAACCA | TGCTTGCTAAATCAGGGAAAA |
| Homo Sapiens | *CEBPA* | AGCTTTCTGGTGTGACTCGG | TATAGGCTGGGCTTCCCCTT |

**Supplemental Table 2: Participant Characteristics for Figure 1C**.

| Physical Characteristics | Non-PAD (N=18) | CLTI (N=38) | *P* value ($\chi^2$ test or *t*-test) |
|---|---|---|---|
| Mean age, y (SD) | 66.7 (10.6) | 64.6 (8.8) | 0.60 |
| Female sex, n (%) | 6 (30.0) | 14 (36.8) | 0.79 |
| Non-White race, n (%) | 4 (22.2) | 8 (21.0) | 0.80 |
| Non-Hispanic, n (%) | 17 (94.4) | 36 (94.7) | 0.55 |
| Disease Characteristics | | | |
| ABI (SD) | 1.01 (0.08) | 0.40 (0.15) | 2.4E-06 |
| Rutherford Stage IV, n (%) | N/A | 19 (50.0) | N/A |
| Rutherford Stage V, n (%) | N/A | 19 (50.0) | N/A |
| Tobacco use | | | |
| Former smoker, n (%) | 10 (55.5) | 27 (71.0) | 0.39 |
| Current smoker, n (%) | 3 (16.6) | 8 (21.0) | 0.69 |
| Medical History | | | |
| Diabetes type I or II, n (%) | 7 (38.8) | 20 (52.6) | 0.49 |
| Hypertension, n (%) | 14 (77.7) | 34 (89.4) | 0.44 |
| Hyperlipidemia, n (%) | 12 (66.6) | 34 (89.4) | 0.08 |
| Coronary artery disease, n (%) | 7 (38.8) | 19 (50.0) | 0.62 |
| Congestive heart failure, n (%) | 5 (27.7) | 9 (23.6) | 0.74 |
| Medication used | | | |
| Aspirin, n (%) | 11 (61.1) | 35 (76.1) | 0.0141 |
| ACE inhibitor, n (%) | 3 (16.6) | 14 (36.8) | 0.22 |
| Angiotensin receptor blocker, n (%) | 3 (16.6) | 6 (15.7) | 0.76 |
| Statin, n (%) | 12 (66.6) | 34 (89.4) | 0.08 |
| Cilostazol, n (%) | 0 (0.0) | 8 (21.0) | 0.09 |
| Beta Blocker, n (%) | 14 (50.0) | 20 (52.6) | 0.13 |
| Anticoagulant, n (%) | 3 (16.6) | 19 (50.0) | 0.0364 |
| Antiplatelet, n (%) | 5 (27.7) | 14 (36.8) | 0.71 |

**Supplemental Table 3: Statistical Reporting of Major Outcomes in mFATGAIN mice.**

| | Control | | | | mFATGAIN | | | |
|---|---|---|---|---|---|---|---|---|
| | Male | | Female | | Male | | Female | |
| **Outcome Variable** | *7dpi* | *28dpi* | *7dpi* | *28dpi* | *7dpi* | *28dpi* | *7dpi* | *28dpi* |
| Total Capillaries/$mm^2$ | 371.8 (65.4) | 411.7 (82.4) | 429.8 (79.2) | 433.6 (118.2) | 406.2 (106.2) | 377.2 (120.0) | 390.6 (86.4) | 418.0 (133.2) |
| $PERILIPIN^+$ area/$mm^2$ | 0.27 (0.13) | 0.56 (0.30) | 0.50 (0.72) | 1.56 (1.10) | 5.67 (1.30) | 5.62 (3.11) | 6.92 (2.48) | 8.76 (2.51) |
| Total Work (Joules) | 5235 (1301) | 7248 (1861) | 3581 (585) | 3411 (878) | 1163 (1190) | 5094 (2244) | 619 (460) | 1656 (829) |

Data presented as mean (SD).

**Supplemental Table 4: Statistical Reporting of Major Outcomes in mFATBLOCK mice.**

| | Control | | | | mFATBLOCK | | | |
|---|---|---|---|---|---|---|---|---|
| | Male | | Female | | Male | | Female | |
| **Outcome Variable** | *7dpi* | *28dpi* | *7dpi* | *28dpi* | *7dpi* | *28dpi* | *7dpi* | *28dpi* |
| Total Capillaries/$mm^2$ | 284.7 (40.2) | 335.3 (88.5) | 337.2 (88.3) | 433.3 (86.5) | 306.8 (54.6) | 377.8 (90.8) | 463.4 (73.7) | 434.0 (87.6) |
| $PERILIPIN^+$ area/$mm^2$ | 1.74 (1.10) | 1.32 (1.180 | 2.34 (1.23) | 2.28 (1.69) | 0.16 (0.11) | 0.04 (0.03) | 0.14 (0.06) | 0.11 (0.07) |
| Total Work (Joules) | 2380 (2235) | 3496 (1922) | 1183 (1259) | 2361 (1190) | 4141 (1651) | 5732 (1168) | 2611 (570) | 4026 (892) |

Data presented as mean (SD).

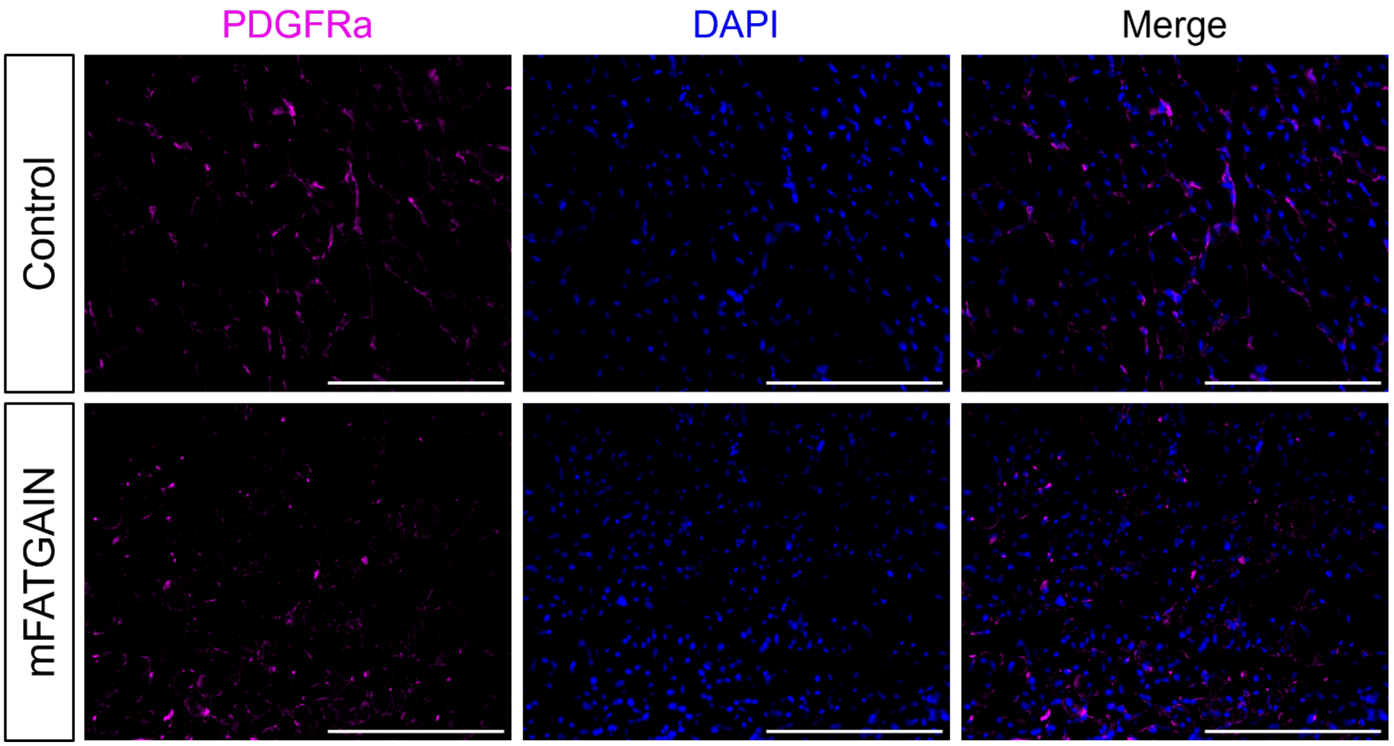


**Supplemental Figure 1. Representative FAP staining in mFATGAIN mice.** Immunofluorescence images of the ischemic gastrocnemius stained with PDGFRa and DAPI in male mice at 28 days post HLI. 20x magnification, scale bars = 200 μm.

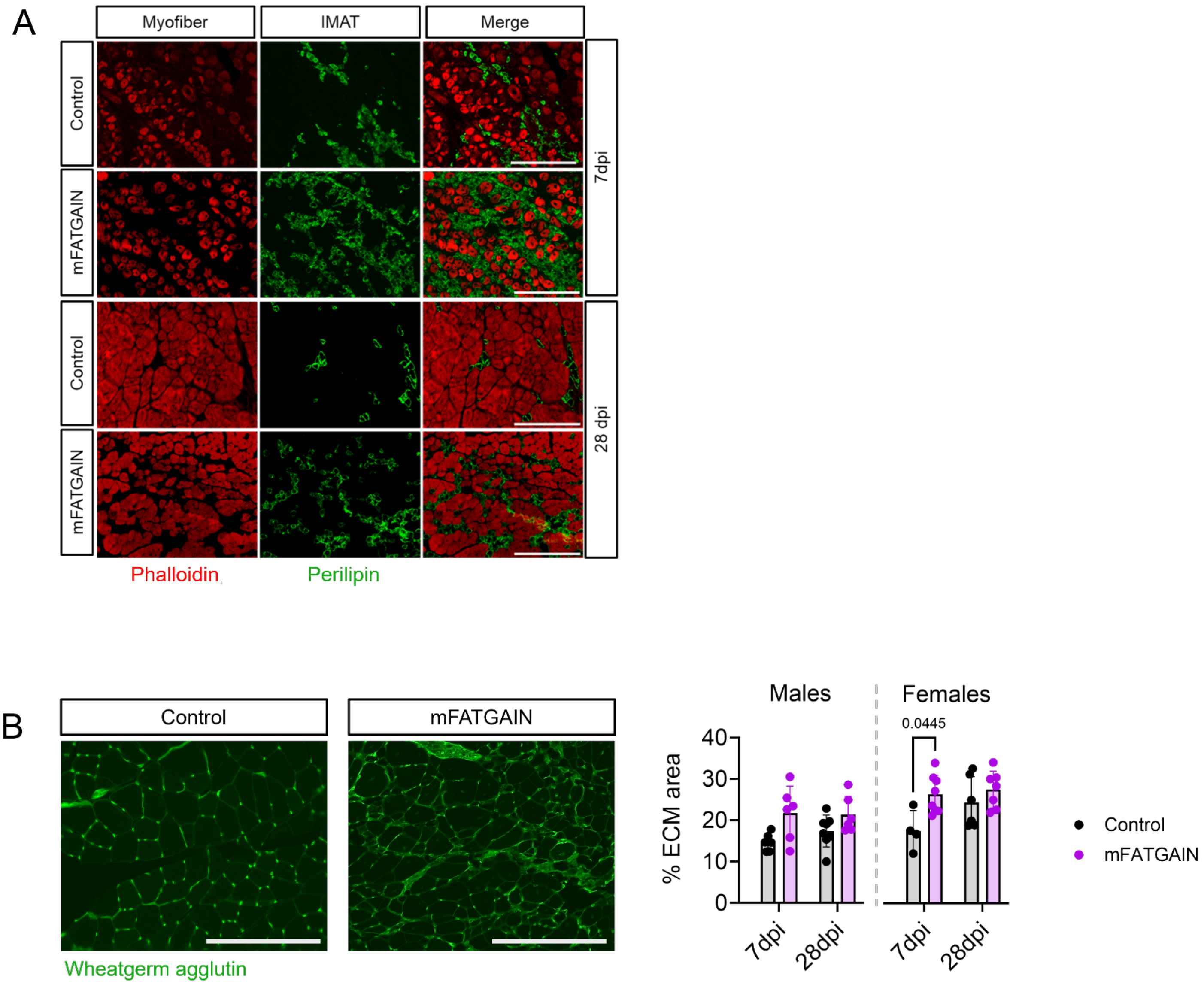


**Supplemental Figure 2. Visualization of IMAT and ECM area in mFATGAIN mice.** (A) Representative images of IMAT (Perilipin) and myofibers (Phalloidin) at 7- and 28-days post HLI. (B) Images and quantification of extracellular matrix expansion reported as percentage of total area. Statistical analyses performed using two-way ANOVA with Tukey post hoc testing for multiple comparisons. Error bars represent the standard deviation. All images at 20x magnification; scale bars = 200 μm.

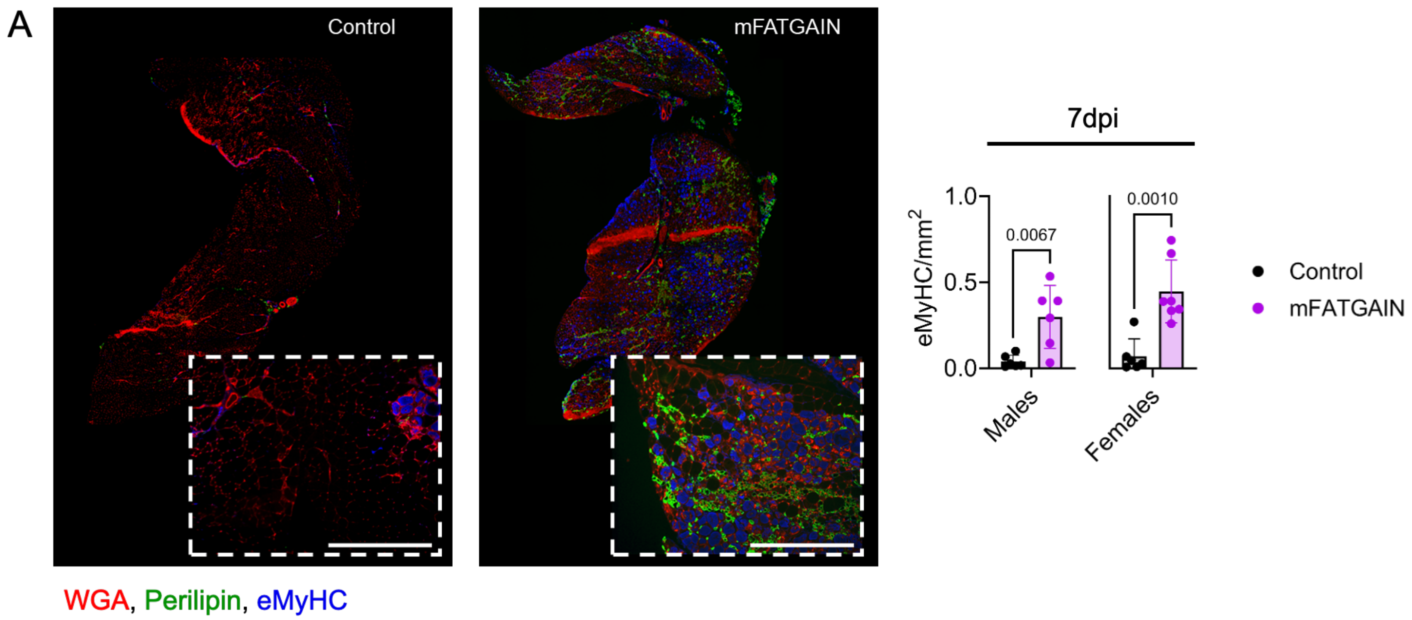


**Supplemental Figure 3. Quantification of embryonic myosin heavy chain positive myofibers in mFATGAIN mice with HLI.** (A) Images and quantification of embryonic myosin heavy chain (eMyHC) staining in the ischemic gastrocnemius 7 days post HLI (n= 6-8 gastrocs/sex/genotype). Data represented as a proportion of total area. Data were analyzed using an unpaired t-test and error bars represent ± SD All images at 10x magnification; scale bars = 400 μm.

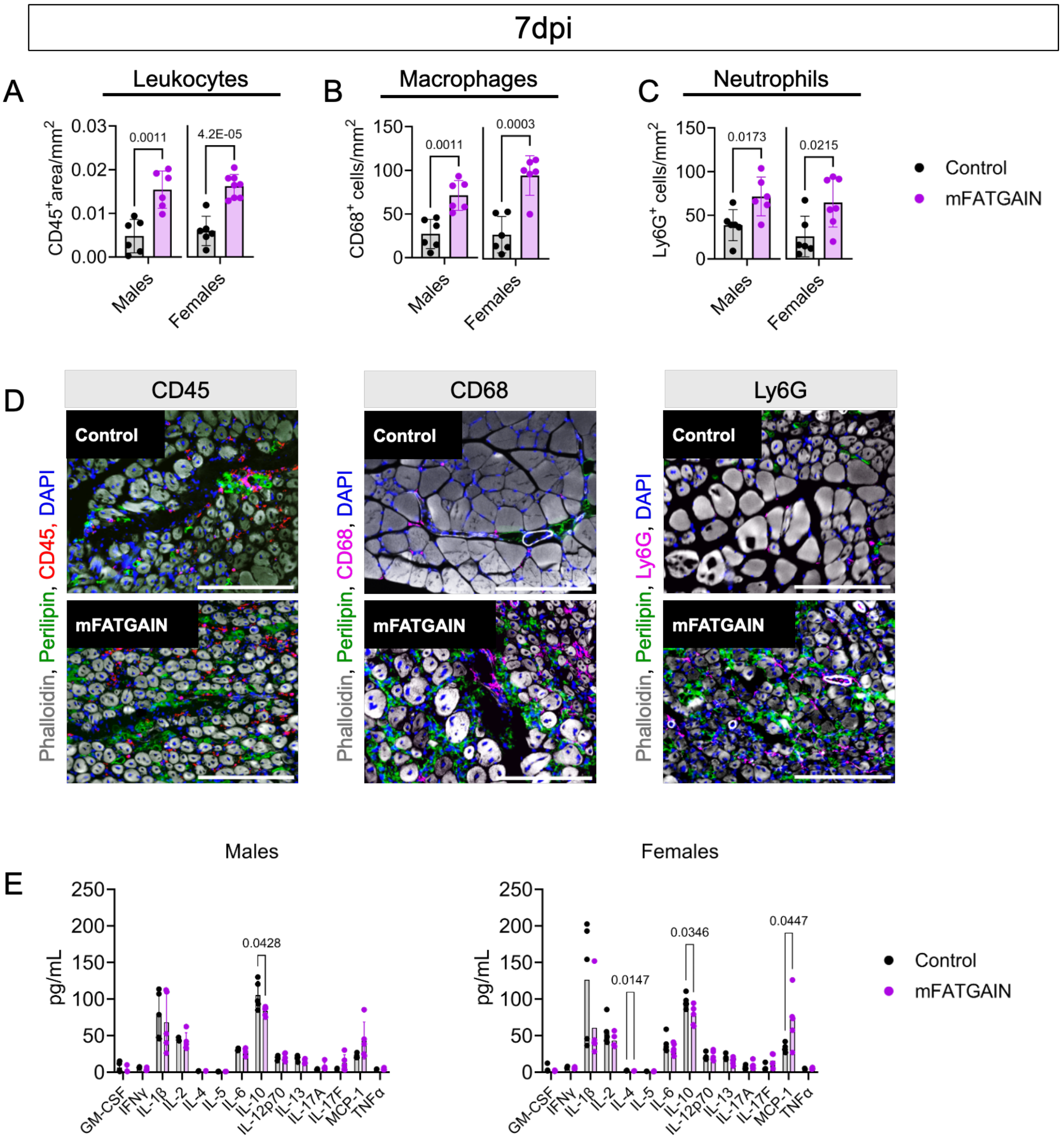


**Supplemental Figure 4. Immune cell abundance and cytokine levels in mFATGAIN mice with HLI.** (A-D) Quantification of immune cells normalized to total area from the ischemic limb 7 days post HLI (n=6-8 /sex/genotype). (A) Leukocytes quantified using CD45 staining, (B) Neutrophils quantified using Ly6G staining, and (C) Macrophages quantified using CD68 staining. (D) Representative images of immune cells at 20x magnification; scale bars = 200 µm. (E) Cytokine analysis of GM-CSF, IFNγ, IL-1β, IL-2, IL-4, IL-5, IL-6, IL-10, IL-12p70, IL-13, IL-17A, IL-17F, MCP-1/CCL2, and TNFα using whole muscle lysate from the ischemic plantaris and soleus 7 days post HLI (n=5/sex/genotype). Data were analyzed using an unpaired t-test and error bars represent ± SD.

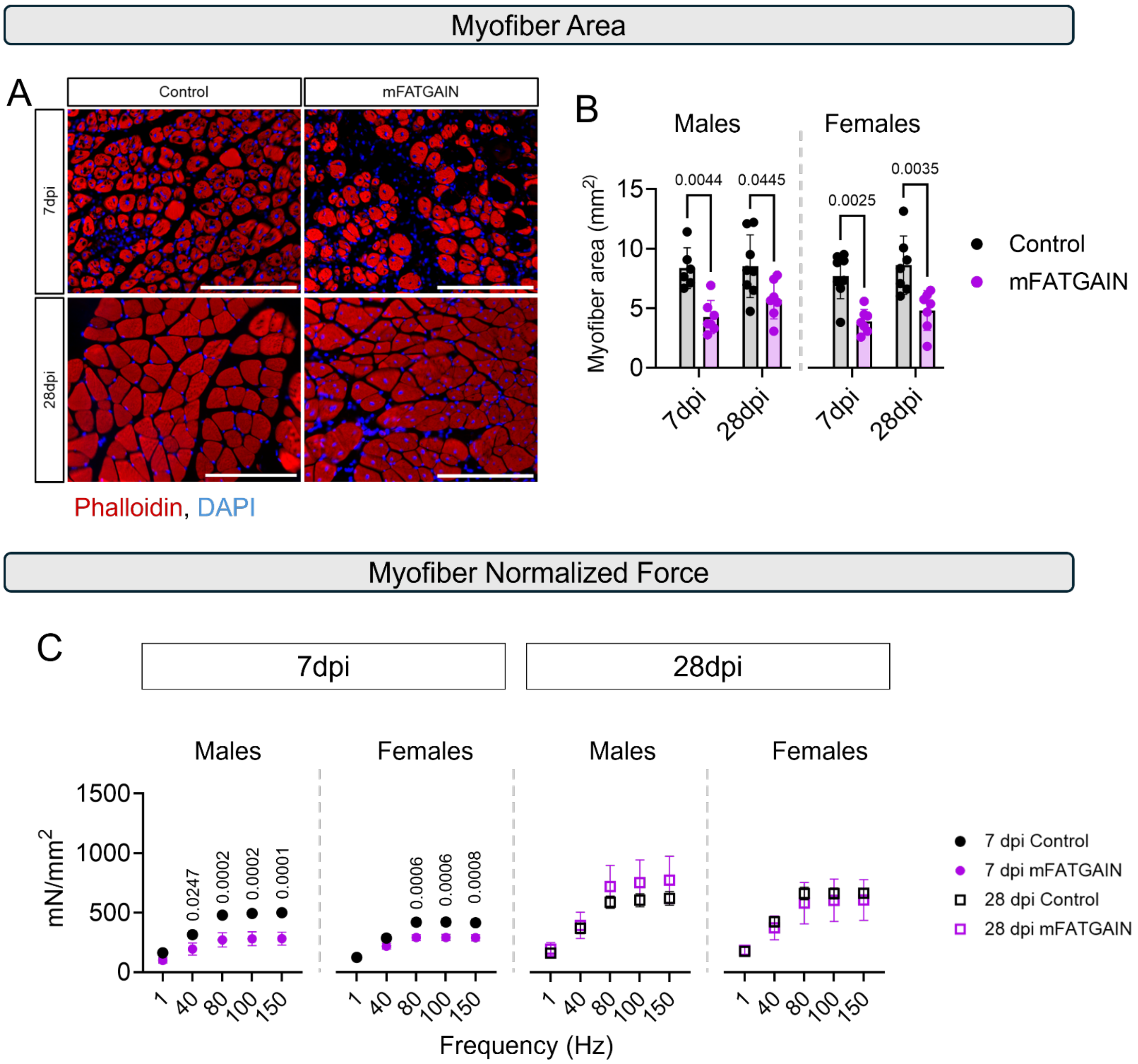


**Supplemental Figure 5. IMAT associated strength deficits are not caused by intrinsic myofilament protein damage.** (A) Representative Phalloidin staining at 20x magnification to visualize myofibers at 7- and 28-days post HLI; scale bars = 200 µm. (B) Total myofiber area quantified. (C) Absolute muscle force values normalized to total myofiber area. (B,C) Males (7dpi; n = 6/genotype. 28dpi; control n = 8, mFATGAIN n = 7) and females (7dpi; n=8 mice/genotype 28dpi; n=7 mice/genotype). All data are represented as mean ± SD.

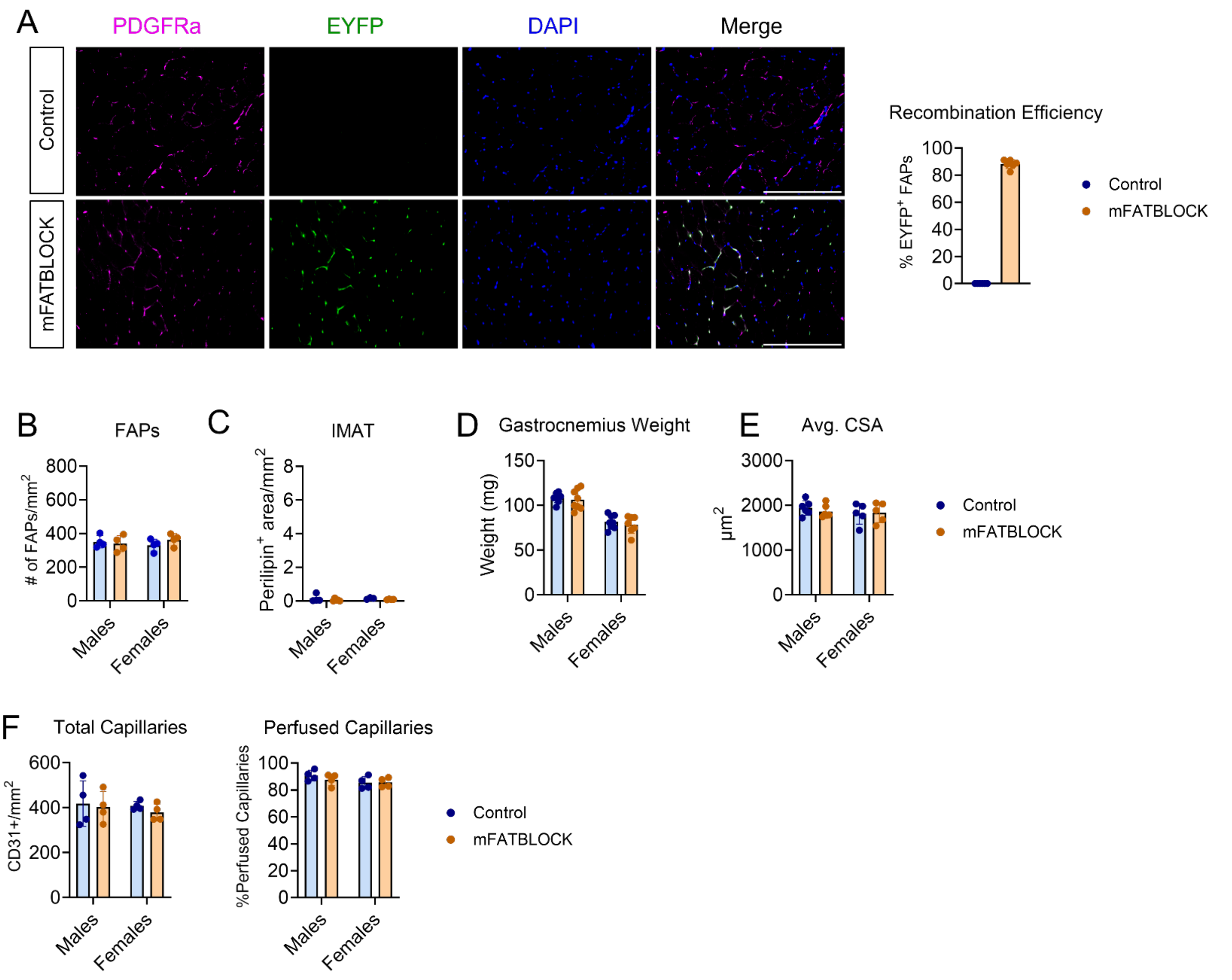


**Supplemental Figure 6. FAP-Specific Deletion of Pparγ does not alter IMAT in non-ischemic muscle.** (A-F) Analysis of non-ischemic gastrocnemius muscle. (A) Detection of EYFP reporter in FAPs and quantifications of the percentage $EYFP^+$ FAPs of FAPs and total FAPs (B). (C) Quantification of IMAT measured with Perilipin staining (n=3-4/genotype/sex). (D) Muscle wet weight (n=7-8/genotype/sex). (E) Average myofiber cross-sectional area (CSA) (males n=6/genotype and females n=5/genotype). (F) Total capillaries probed with anti-CD31 antibody and normalized to area. Perfused capillaries labeled with retro-orbitally injected isolectin and reported as a percentage of total capillaries (n=4/genotype/sex). Images are 20x magnification, scale bars = 200 µm. All data are represented as mean ± SD.

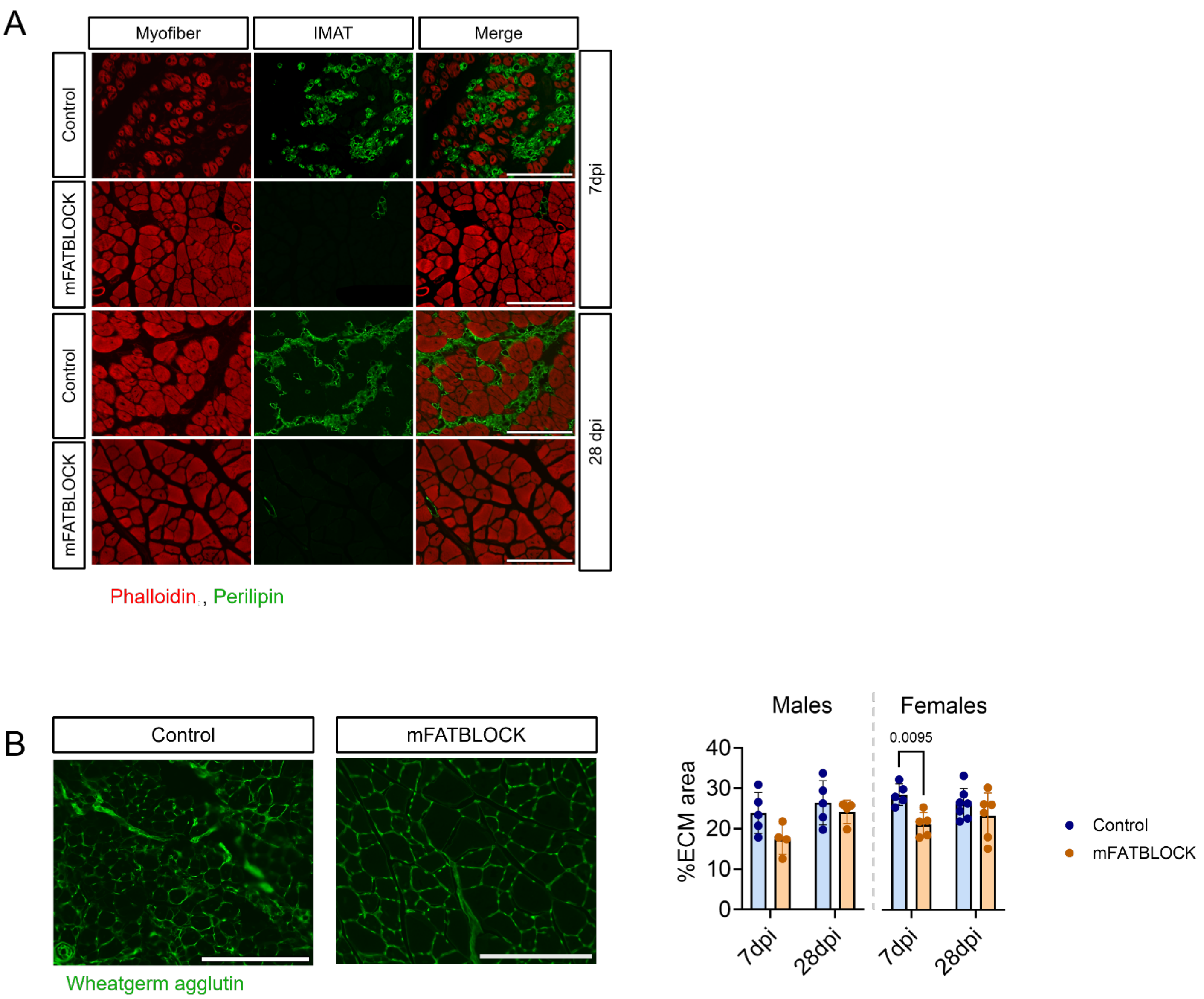


**Supplemental Figure 7. Visualization of IMAT and ECM area in mFATBLOCK mice.** (A) Representative images of IMAT (Perilipin) and myofibers (Phalloidin) at 7- and 28-days post HLI. (B) Images and quantification of extracellular matrix expansion reported as percentage of total area. (n=4-6 /genotype/sex/timepoint). Statistical analyses performed using two-way ANOVA with Tukey post hoc testing for multiple comparisons. Error bars represent the standard deviation. All images at 20x magnification; scale bars = 200 µm.

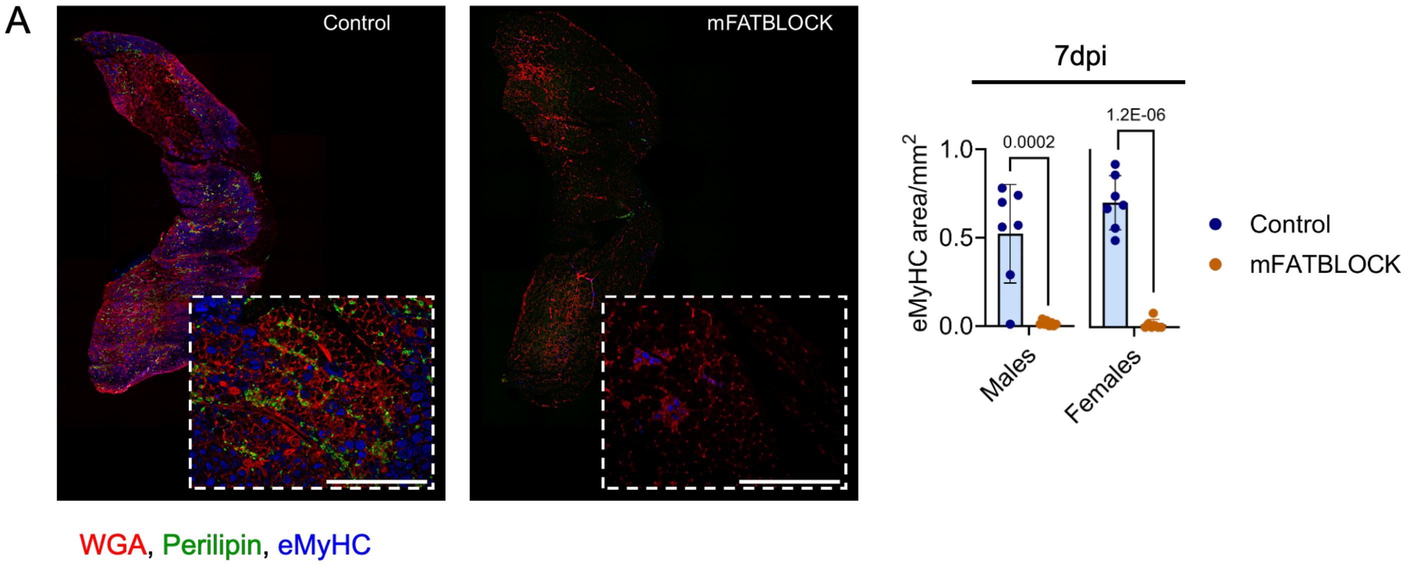


**Supplemental Figure 8. Quantification of embryonic myosin heavy chain positive fibers in mFATBLOCK mice with HLI.** (A) Representative images and quantification of embryonic myosin heavy chain (eMyHC) staining in the ischemic gastrocnemius 7 days post HLI (n= 7-8 sex/genotype). Data represented as a proportion of total area. Data were analyzed using an unpaired t-test and error bars represent ± SD All images at 10x magnification; scale bars = 400 μm.

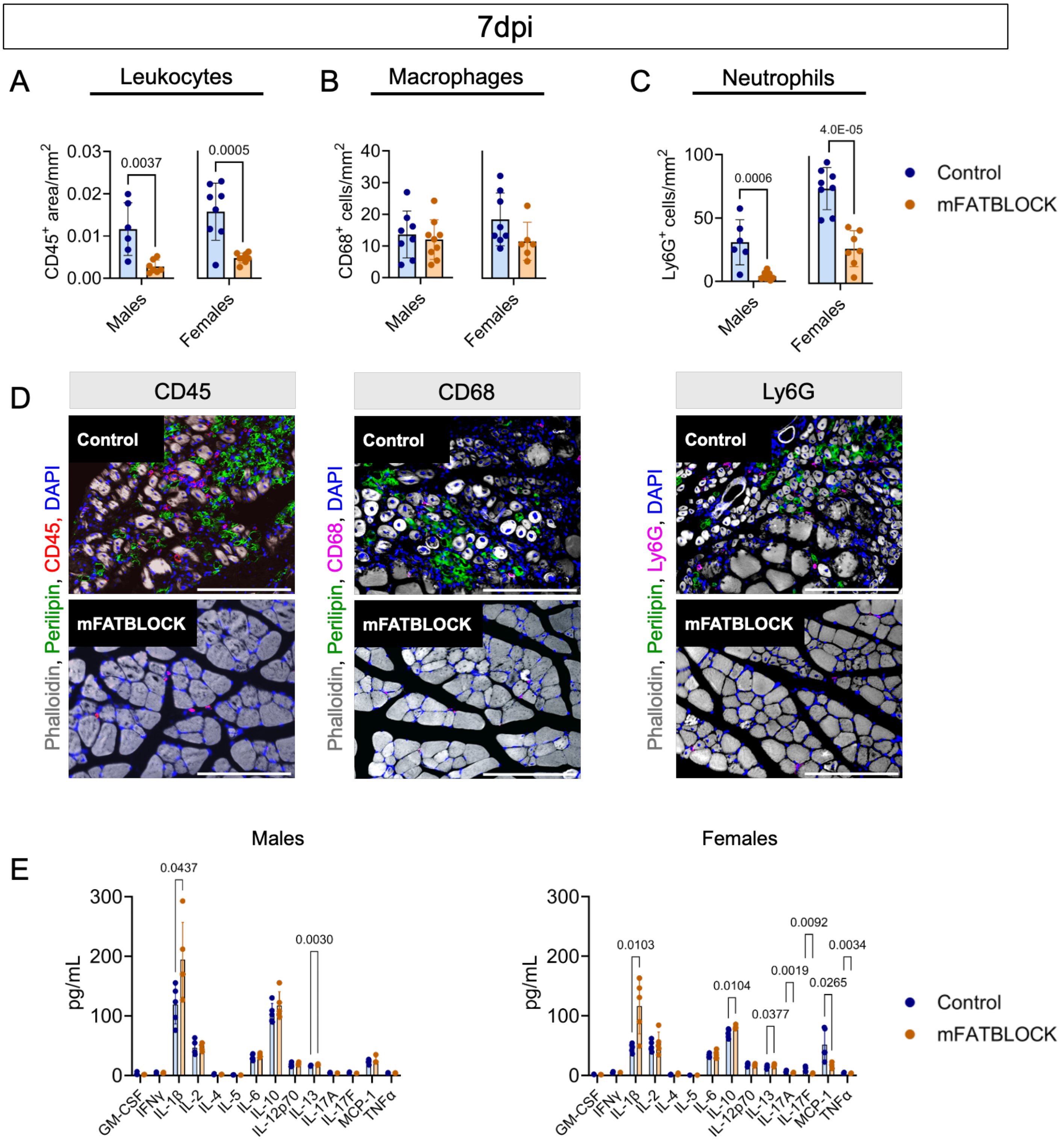


**Supplemental Figure 9. Immune cell abundance and cytokine levels in mFATBLOCK mice with HLI.** (A-D) Quantification of immune cells normalized to total area from the ischemic limb 7 days post HLI (n=7-8 /sex/genotype). (A) Leukocytes quantified using CD45 staining, (B) Neutrophils quantified using Ly6G staining, and (C) Macrophages quantified using CD68 staining. (D) Representative images of immune cells at 20x magnification; scale bars = 200 μm. (E) Cytokine analysis of GM-CSF, IFNγ, IL-1β, IL-2, IL-4, IL-5, IL-6, IL-10, IL-12p70, IL-13, IL-17A, IL-17F, MCP-1/CCL2, and TNFα using whole muscle lysate from the ischemic plantaris and soleus 7 days post HLI (n=5/sex/genotype). Data were analyzed using an unpaired t-test and error bars represent ± SD.

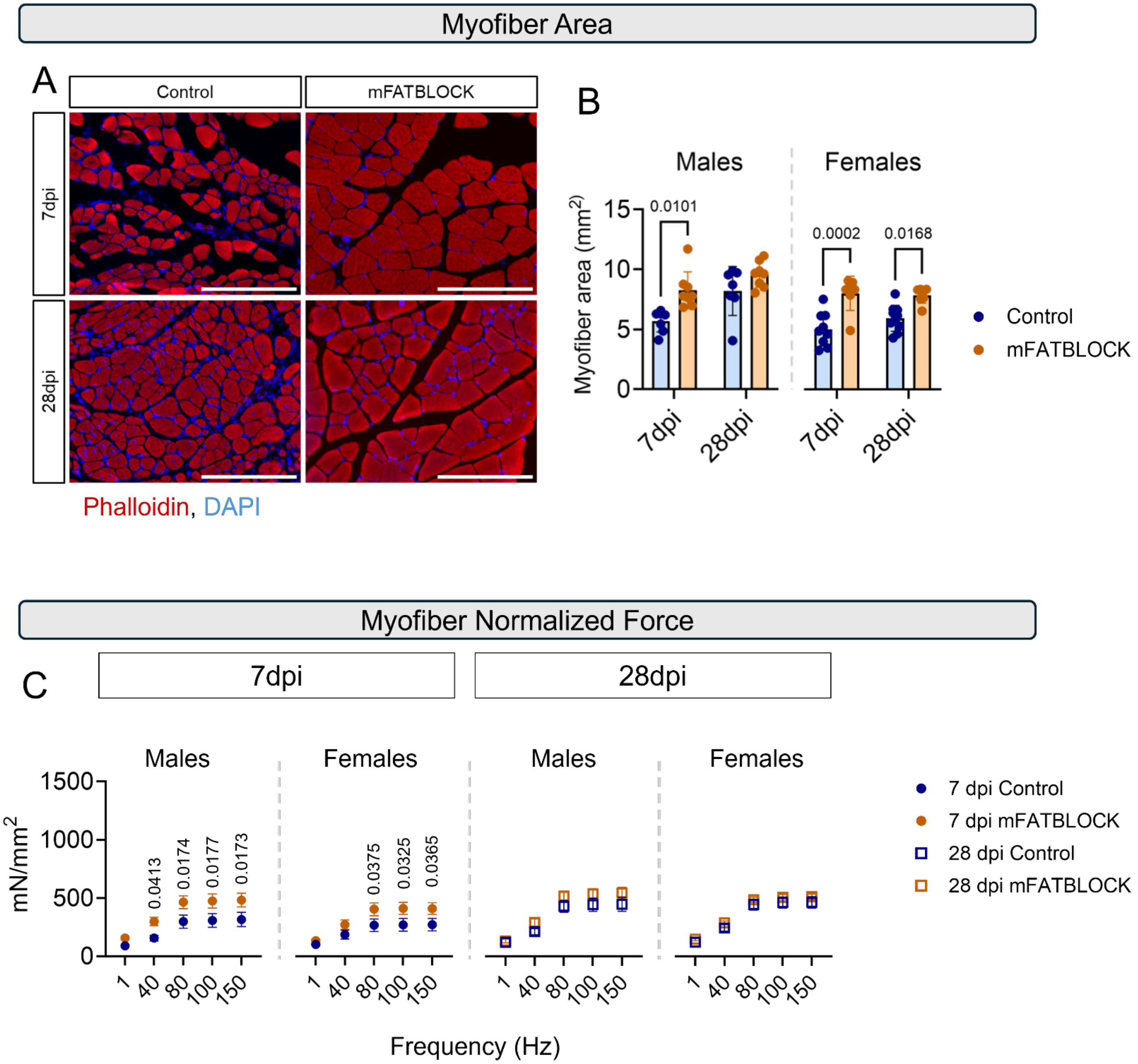


**Supplemental Figure 10. Ablation of IMAT accelerates recovery of myofilament contractile function.** (A) Representative Phalloidin staining at 20x magnification to visualize myofibers at 7- and 28-days post HLI; scale bars = 200 μm. (B) Total myofiber area quantified. (C) Absolute muscle force values normalized to total myofiber area. (n= 7-9/genotype/sex/timepoint).

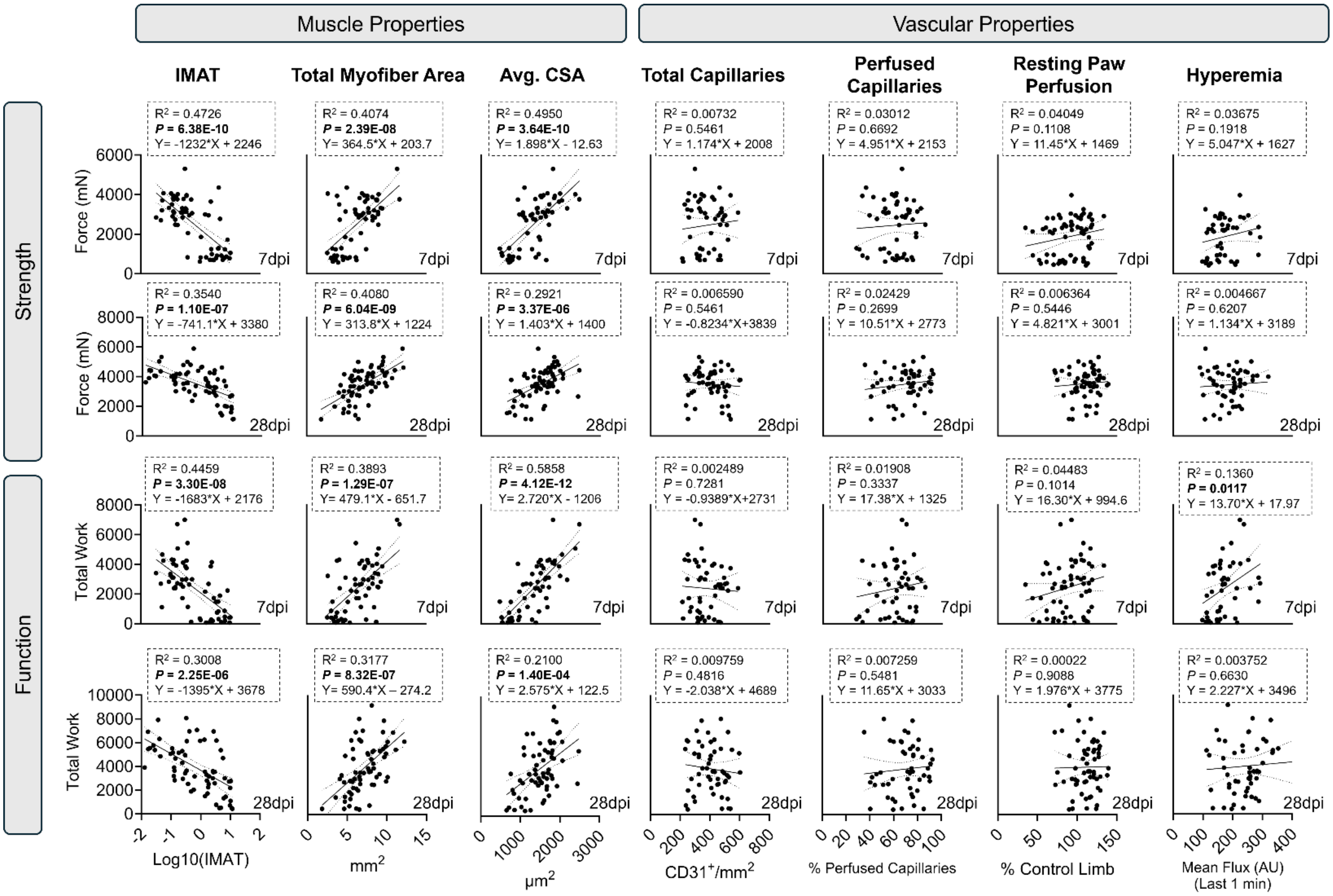


**Supplemental Figure 11. Traditional outcome measures are not correlated with improvements in muscle function in mice with HLI.** Linear regression analysis of absolute force generated at 80 Hz stimulation versus IMAT, total myofiber area, average cross-sectional area (CSA), total capillaries, perfused capillaries, resting paw perfusion, and hyperemia and total work versus IMAT, total myofiber area, average cross-sectional area (CSA), total capillaries, perfused capillaries, resting paw perfusion, and hyperemia in mice 7- or 28-days post HLI. Data were pooled from all mice across study and analyzed using a simple linear regression and Pearson correlation test.

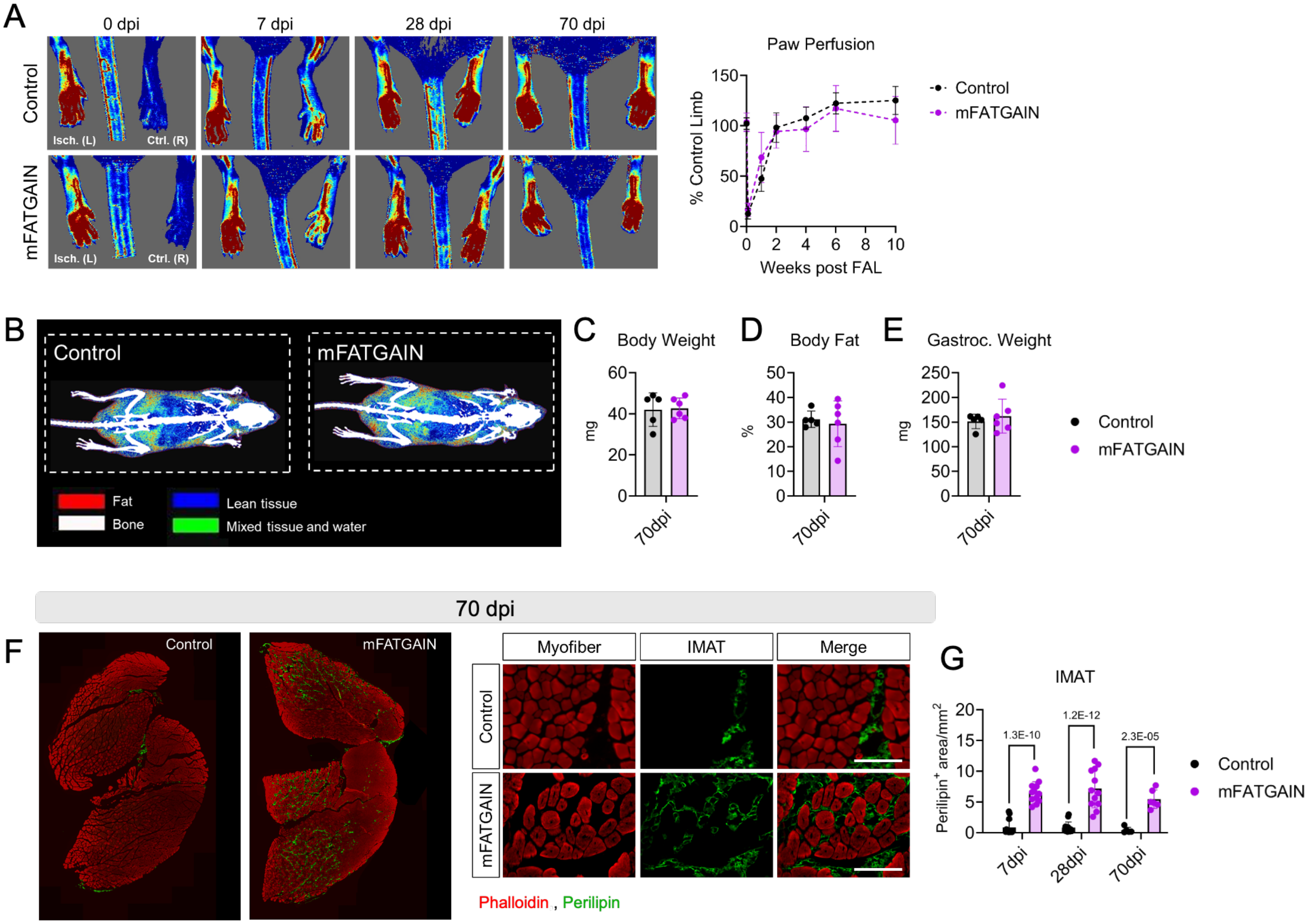


**Supplemental Figure 12. Increased IMAT accumulation persists in mFATGAIN mice with HLI following a 70-day recovery period:** (A) Resting paw perfusion measured with Laser Doppler imaging throughout the 10-week recovery period and analyzed using a mixed-effects model (n=5/genotype/timepoint). (B-F) Data collected at 70 days post HLI. (B) Representative body composition scans using Dual-energy X-ray absorptiometry (DEXA). (C) Body weight and (D) total body fat. (E) Wet weight of the gastrocnemius of the ischemic limb. (F) Representative images at 20x magnification of IMAT (Perilipin) and myofibers (Phalloidin); scale bars = 200 μm. (G) Quantification of IMAT in combined male and female mFATGAIN mice at 7, 28, and 70-days post HLI. (C-E) Statistical analysis was performed using a Mann-Whitney U test (n= 5/genotype). (G) Data were analyzed using a two-way ANOVA with Tukey's post hoc testing for multiple comparisons when significant interactions were detected. (7dpi n=12-13/genotype; 28 dpi n=12-14/genotype; 70dpi n=5/genotype). All data are presented as mean ± SD.

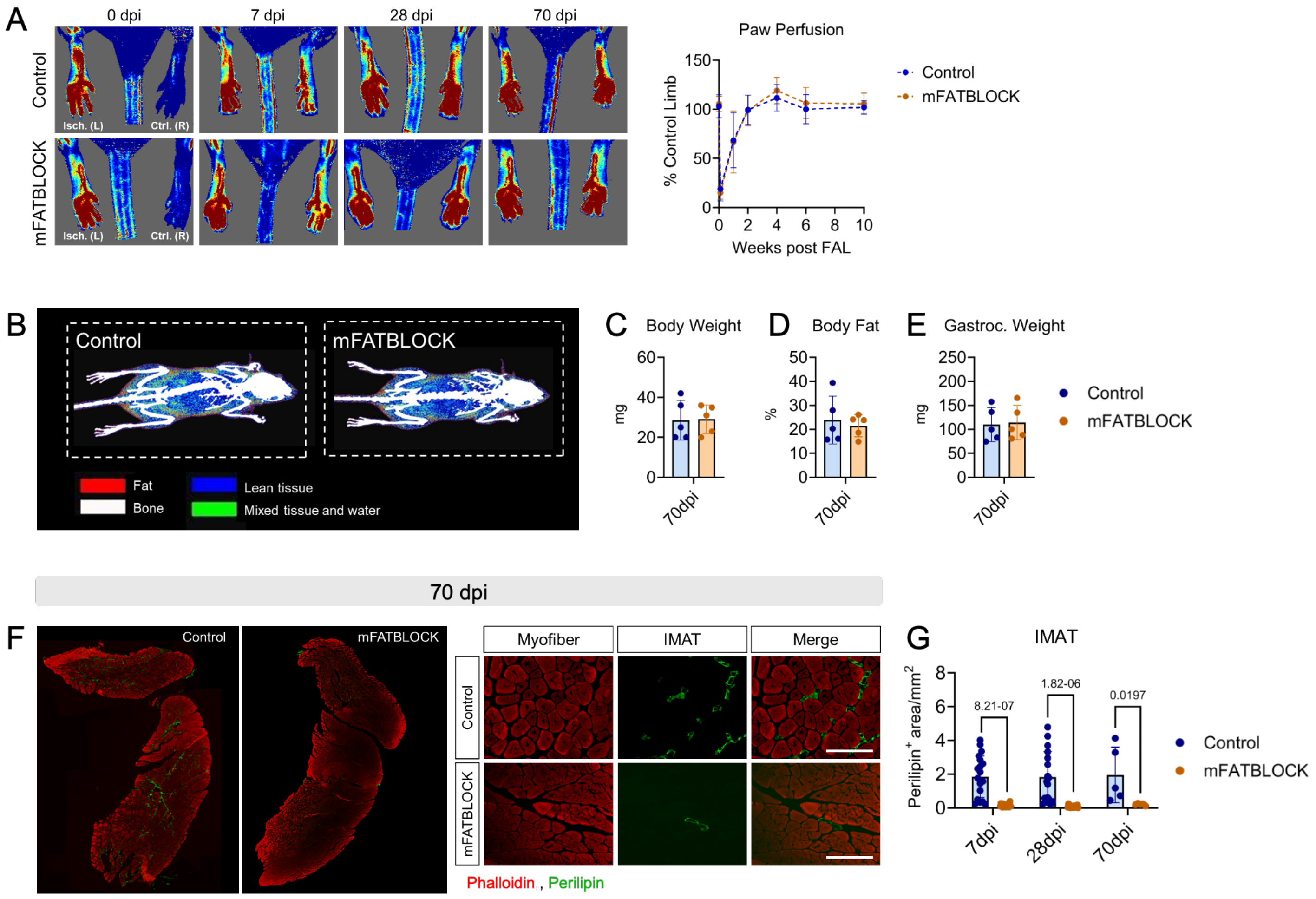


**Supplemental Figure 12. IMAT remains suppressed in mFATBLOCK mice with HLI following a 70-day recovery period:** (A) Resting paw perfusion measured with Laser Doppler imaging throughout the 10-week recovery period and analyzed using a mixed-effects model (n=5/genotype/timepoint). (B-F) Data collected at 70 days post HLI. (B) Representative body composition scans using Dual-energy X-ray absorptiometry (DEXA). (C) Body weight and (D) total body fat. (E) Wet weight of the gastrocnemius of the ischemic limb. (F) Representative images at 20x magnification of IMAT (Perilipin) and myofibers (Phalloidin); scale bars = 200 µm. (G) Quantification of IMAT in combined male and female mFATBLOCK mice at 7, 28, and 70-days post HLI. (C-E) Statistical analysis was performed using a Mann-Whitney U test (n= 5/genotype). (G) Data were analyzed using a two-way ANOVA with Tukey's post hoc testing for multiple comparisons when significant interactions were detected. (7dpi n=19-20/genotype; 28 dpi n=17/genotype; 70dpi n=5/genotype). All data are presented as mean ± SD.

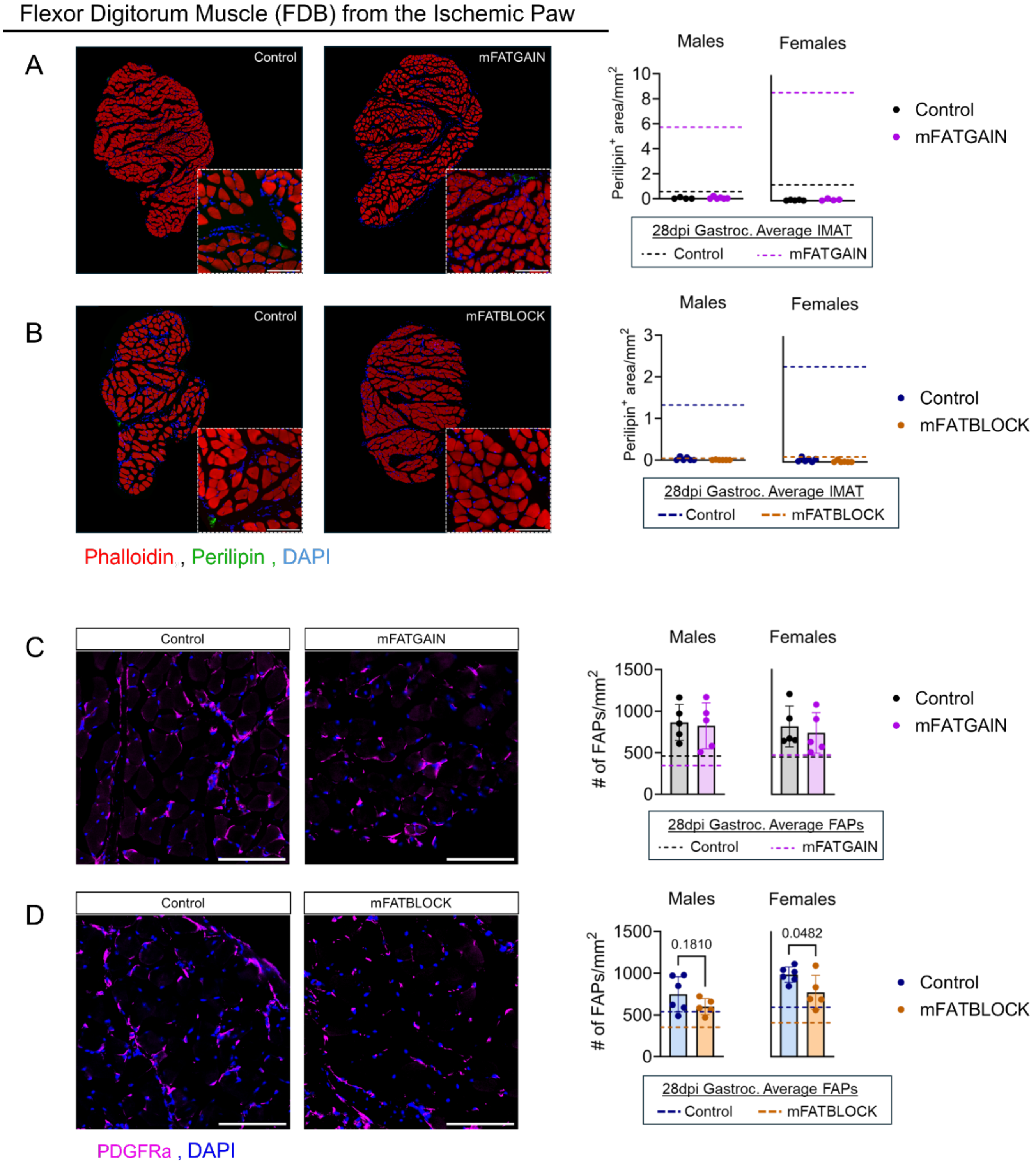


**Supplemental Figure 14. Flexor Digitorum Brevis (FDB) muscle does not form IMAT following ischemia.** Representative images and quantification of IMAT (Perilipin$^+$ area) in the FDB muscle from the ischemic paw of mFATGAIN (A) and mFATBLOCK (B) mice, as well as their respective littermate controls, 28-days post hindlimb ischemia. Representative images and quantification of FAP abundance (PDGFRA$^+$ cells) in the FDB muscle from the ischemic paw of mFATGAIN (C) and mFATBLOCK (D) mice, as well as their respective littermate controls, 28-days post hindlimb ischemia. Images at 20x magnification, scale bars = 100μm. Data were analyzed using an unpaired t-test and error bars represent ± SD (n=5-6 FDBs/sex/genotype). Dashed lines represent the mean values of the ischemic gastrocnemius muscles from the same mice for comparison.

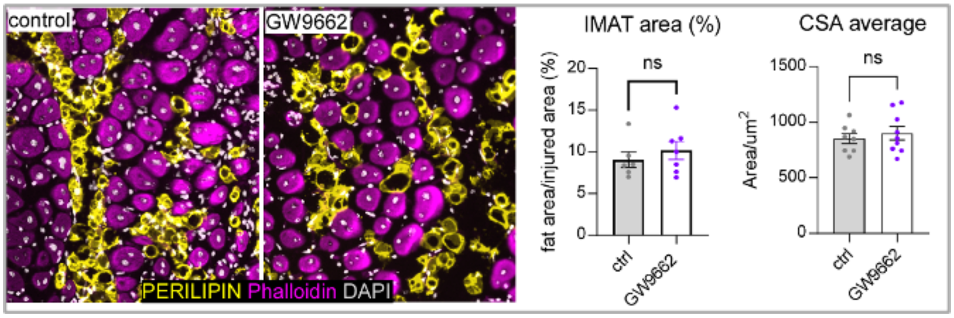


**Supplemental Figure 15. Pparg antagonism fails to repress IMAT formation in non-ischemic muscle injury.** Representative images and quantification of IMAT (Perilipin$^+$ area) in wildtype 129Sv/J mice treated with placebo (ctrl) or GW9662 following a glycerol injection to cause injury of the tibialis anterior muscle. Quantification of the IMAT area and mean myofiber cross sectional area (CSA) indicated no repression of IMAT formation. Data were analyzed using an unpaired t-test and error bars represent ± SEM (n=6-9 male mice per group).
